# Low-Cost Screening of Algae for Extreme Tolerance to pH, Temperature, Salinity, and Light

**DOI:** 10.1101/2025.05.12.653396

**Authors:** Barbara Saucedo, Kalisa Kang, Lauren May, Abhishek Gupta, Évellin do Espirito Santo, Stephen Mayfield, João Vitor Dutra Molino

## Abstract

Bioprospecting algae strains with tolerance to extreme conditions such as pH, temperature, salinity, and light is crucial for advancing biotechnology and environmental applications. However, traditional screening methods often involve significant costs and labor, restricting their accessibility and practical use. In this study, we developed and validated low-cost, high-throughput screening techniques, predominantly employing agar plates and liquid culture assays, to effectively differentiate tolerance levels among various algae strains. The methodologies were optimized using the model microalga *Chlamydomonas reinhardtii* and its closely related species *Chlamydomonas incerta* and the recently discovered extremophilic *Chlamydomonas pacifica*. We systematically evaluated the algae for tolerance to extremes by establishing precise gradients of pH (acidic to alkaline conditions), salinity (0 to 5 M NaCl), temperature (34–42°C), and light intensity (40 to 2977 μE·m⁻²·s⁻¹). Our results demonstrated that these cost-effective, agar plate-based methods effectively distinguished algae strains exhibiting superior tolerance to extreme environmental conditions. These screening techniques not only provided clear differentiation among the closely related strains but also delivered reproducible outcomes suitable for scaling up to larger bioprospecting efforts. Furthermore, the affordability and simplicity of these methods facilitate their implementation in resource-limited laboratories, thereby broadening participation in algae bioprospecting endeavors. This study highlights the potential of low-cost, accessible screening techniques to significantly enhance the discovery and characterization of algal strains with extreme traits. Ultimately, these methods support the development of robust algae-based resources, driving innovation in diverse industrial processes and environmental solutions.

**Graphical Abstract:** 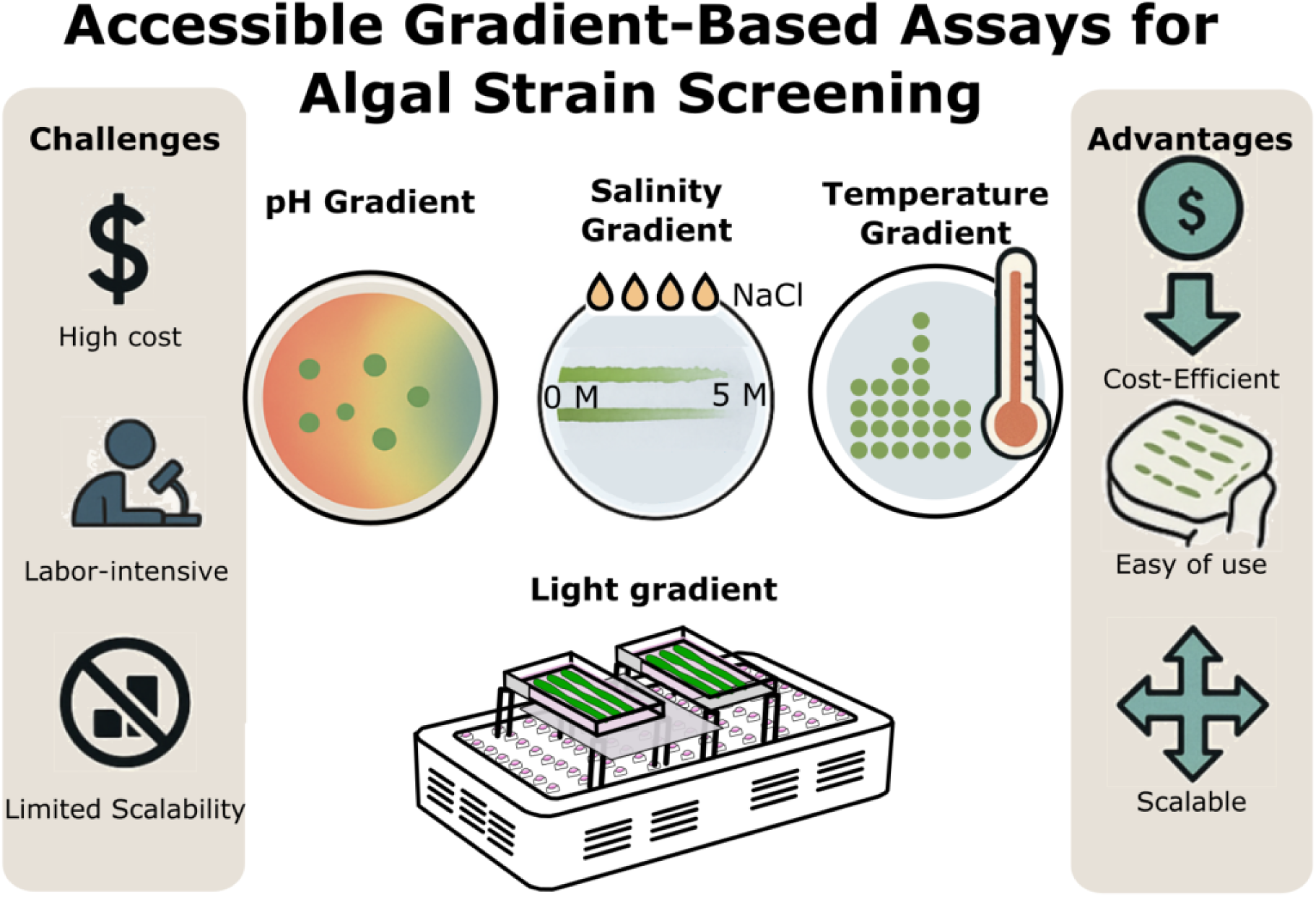

## Introduction

Microalgae are a diverse group of photosynthetic microorganisms that flourish in numerous environments, from freshwater and oceans to soils and even extreme habitats (1). They exhibit remarkable metabolic flexibility and adaptability. Many species can grow photoautotrophically using only sunlight, CO₂ and inorganic nutrients, yet can also adjust their metabolism under suboptimal conditions or adopt mixotrophic growth when organic carbon is available (2,3). This versatility, coupled with rapid biomass accumulation and efficient CO₂ fixation, has positioned microalgae as an attractive platform for sustainable production of fuels, foods and other bioproducts (4). In addition, their capacity to remove excess nutrients and heavy metals (4–6) from wastewater along with their superior CO_2_ capture rates compared to terrestrial plants (7) is appealing to biotechnology and environmental solutions.

Translating microalgal potential into large-scale cultivation is challenging, as many strains cannot withstand the suboptimal or fluctuating conditions encountered in industrial systems. Selecting or developing strains with robust tolerance to stress factors is therefore a critical priority (8,9). Utilizing such stress-tolerant strains (or genetically enhancing tolerance in existing ones) is key to improving reliability and yields in mass culture. Environmental factors like pH, salinity, temperature, and light fundamentally impact algal physiology and thus must be considered in strain selection. Algae cultivated in outdoor raceway ponds or large photobioreactors are constantly subjected to fluctuating environmental conditions (10). Such fluctuations can significantly inhibit growth or induce stress responses, reducing overall productivity (11). Microalgal strains exhibit considerable variability in their tolerance to environmental stressors. Therefore, identifying strains capable of withstanding industrial conditions, such as high light intensity, large daytime temperature swings, overnight cooling, pH shifts, and varying salinity levels, is critical for the successful implementation of large-scale biotechnology processes.

Conventional methods for evaluating algal stress tolerance tend to be laborious, costly, and low-throughput. Strains are tested one at a time in flasks or bioreactors under a specific stress, measuring growth rates or lipid accumulation – a process that does not scale well to hundreds of strains or the wide range of stress gradients that could be explored. The workflow is resource-intensive and time-consuming (12). Moreover, many advanced screening methods — such as flow cytometry (13), high-content imaging (14), or automated turbidostats (15) — rely on specialized instruments that are often inaccessible to labs without substantial infrastructure. This poses a particular barrier for exploring non-model microalgae, which are the vast majority of microalgal species, partly because there are no simple, high-throughput ways to phenotype them for desirable traits (16). While multi-well plate assays and other miniaturized techniques have been introduced to improve throughput (16), these still require expensive plate readers or may not adequately mimic the stress conditions of interest. In short, traditional screening approaches – being costly, time-consuming, and at times low-throughput – fall short of the needs presented by the vast diversity of algae and the range of stresses relevant to industry. There is a need for novel screening methodologies that are scalable (able to handle many strains/conditions at once), high-throughput (generate data rapidly), and accessible (deployable in any laboratory at low cost).

To date, finding stress-tolerant strains has been somewhat ad hoc – relying on either chance isolation from extreme environments or labor-intensive one-by-one testing. A more systematic approach is required to accelerate this discovery and to enable simultaneous selection for multiple stress tolerances. An ideal screening tool would allow researchers to expose many strains (or genetic variants) to a gradient of stress conditions in parallel, in a cost-effective and reproducible manner, so that only the hardiest performers are advanced.

One promising concept is the use of gradient-based plate assays for stress tolerance screening. In microbiology, gradient plate techniques (where a solid agar plate contains a spatial gradient of a stressor, such as an antibiotic or toxin) have long been used for selecting resistant mutants or assessing tolerance thresholds. Such methods are simple, low-cost, and scalable, requiring no specialized equipment beyond basic culturing supplies (17). We propose adapting this approach to microalgae: for example, pouring agar plates that establish a continuous pH gradient from one end to the other, or a salinity or temperature gradient across a multi-well plate.

In this study, we address the need for improved phenotypic screening by introducing a suite of four simple and scalable assays to evaluate microalgal tolerance to pH, salinity, temperature, and high light stress. These assays are designed with accessibility and broad applicability in mind: they employ standard laboratory materials (such as Petri dishes, agar media, and multi-well plates) and do not require sophisticated instruments, making them suitable for virtually any laboratory setting. Each assay establishes a controlled gradient or range of stress conditions and incorporates a strategy for selective recovery of tolerant cells, aligning with the concepts outlined above. We demonstrate application of these methods using three distinct algal strains - including both well-known laboratory species *Chlamydomonas reinhardtii* (Cre) and *Chlamydomonas incerta* (Cin), but also an extremophilic algae *Chlamydomonas pacifica* (Cpa) (9), to showcase their versatility. Key performance metrics — such as growth, survival, or photo-physiological responses—are measured across the stress gradients to determine tolerance thresholds for each strain.

## Results

We evaluated all three strains simultaneously and leveraged recombinant strains carrying distinct antibiotic resistance markers (bleomycin and hygromycin) to assess the ability of the screening methods to distinguish between strains under the four tested stress conditions.

Among the strains, Cin exhibited the greatest sensitivity to salt, pH, and high temperature. Next, Cre demonstrated intermediate sensitivity to these environmental stressors. Lastly, Cpa showed the greatest tolerance across all stressors. Cin exhibited a moderate light tolerance in comparison to *Cre,* The detailed methods and corresponding results are presented below.

### pH Gradient plate

#### Formation and Stabilization of a pH Gradient

The formation and stabilization of a pH gradient on High Salt Media (HSM) media with phosphate as a buffer system were monitored over time, as shown in **Figure 1**. A fixed volume of 120 uL of 10 M KOH solution (blue dots) was applied to one corner of the plate, while a volume of 60 uL of 3.3 M phosphoric acid solution (red dots) was added to the opposite corner (panels 1–2). After allowing diffusion and drying, 1 mL of Universal pH Indicator Solution was uniformly applied to visualize the pH distribution across the plate surface. The Fisher Chemical™ Universal pH Indicator System was used to visualize pH gradients, producing a distinct color spectrum ranging from red (<pH 4.0, acidic) to dark blue (>pH 10.0, strongly basic). This color scale identifies pH transitions, with red to yellow indicating acidic conditions, green representing near-neutral pH (around 7.0), and cyan to blue signifying progressively basic environments (**Supplementary Figure 1**). At day 0 (**0d**, panel 3), no visible pH gradient was observed immediately after indicator application, as shown by minimal color variation across the plate. By day 1 (**1d**, panel 4), a clear pH gradient emerged, with visible transitions from acidic (red) to basic (blue) regions. The gradient continued to develop, displaying a smooth color transition from acidic to neutral (green) and basic regions by day 3 (**3d**, panel 5). At this stage, the pH gradient appeared stable and well-defined. We measured the progression of the acidic front (pH ∼4.0–5.0) over time in a time lapse video (**[Supplementary Video 1](https://www.youtube.com/watch?v=4yDHsPnsOm8&feature=youtu.be)**), as shown in **Supplementary Figure 2**, which reveals rapid expansion within the first 3 days, followed by a long period of stability between days 3 and 17 (∼14 days) at ∼60 mm from the plate edge, which is more than sufficient for most algae cells to grow and be selected by pH, before a gradual decline start to be observed after 12 days due to diffusion dynamics or buffer interactions. The results highlight a time-dependent progression of gradient formation, with stabilization occurring at day 3 and remaining stable for 14 days (With in 10% variation). This demonstrates the effective diffusion and persistence of the pH gradient across phosphate media (HSM, nitrate), as visualized using the pH indicator.

**Figure 1:**
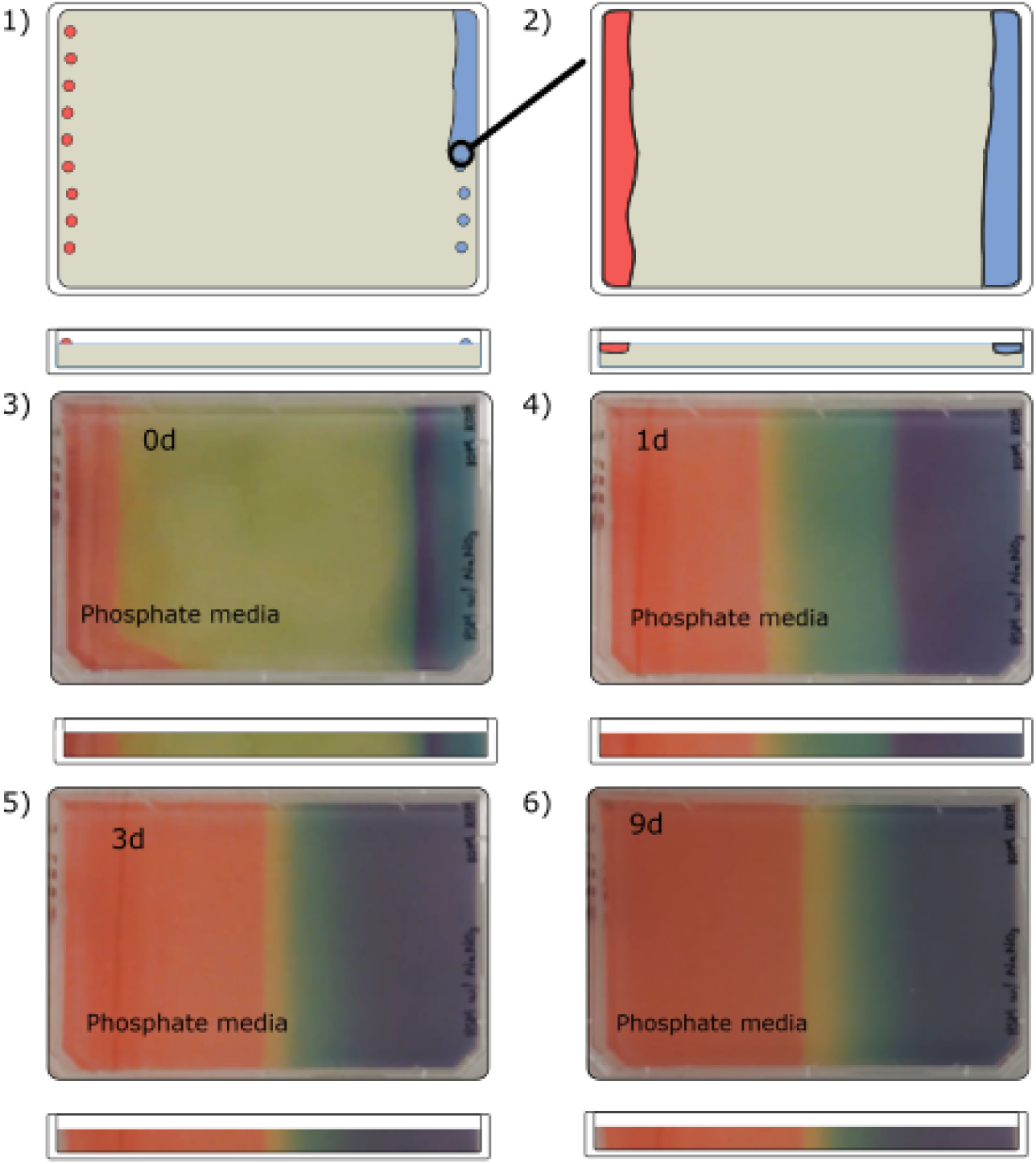
Schematic and visual demonstration of pH gradient formation on HSM nitrate media. (*1–2*) Schematic diagrams illustrating the experimental setup for pH gradient formation on phosphate media. A fixed volume (120 uL) of 10 M KOH solution was applied to one corner of the plate (blue dots), while a fixed volume (60 uL) of 3.3 M phosphoric acid was added to the opposite corner (red dots). After the acid and base solutions were allowed to diffuse and dry, 1 mL of Fisher Chemical™ Universal pH Indicator Solution (Catalog number: SI60-500) was uniformly spread over the entire surface of the plate to visualize the pH gradient. The gradient bar below each panel represents the transition from acidic (red) to basic (blue) regions across the plate. (*3–6*) Time-lapse images showing the development and stabilization of the pH gradient in phosphate media plates over time. (*3*) At 0 days (0 d), the plate shows minimal visible color changes immediately after indicator application. (*4*) After 1 day (1 d), a clear pH gradient becomes visible, with the indicator color representing the pH variation across the plate. (*5*) By 3 days (3 d), the gradient stabilizes, displaying distinct and continuous color transitions from acidic (red) to basic (blue). (*6*) After 9 days (9 d), the gradient remains stable, with no noticeable changes in the distribution of pH across the plate. Gradient bars below each image highlight the corresponding pH-dependent color spectrum at each time point.

#### Buffer-Dependent Tuning of pH Gradients

The development of pH gradients was evaluated on plates containing phosphate media (HSM), citrate buffer, and a combination of phosphate media (HSM) with citrate buffer (**[Supplementary Video 1](https://www.youtube.com/watch?v=4yDHsPnsOm8&feature=youtu.be)**). Clear differences were observed in the formation and extent of the pH gradients across the three buffer systems. In the **phosphate media (HSM)** plate, a well-defined and stable pH gradient was observed, with a smooth transition from acidic regions (red) through neutral (green) to basic regions (purple). This demonstrates effective buffering and diffusion properties within the phosphate media system. The **citrate buffer** plate also exhibited a distinct pH gradient, with a different distribution across the plate, with other regions retaining specific pHs. This suggests the capacity of this approach to tailor the pH gradient with more spread pH variation in different pH ranges, depending on the buffer system used. For the **phosphate media (HSM) + citrate buffer** plate, the pH gradient appeared broader and more gradual, with a clear transition spanning red (acidic), yellow, green (neutral), and purple (basic) regions. This indicates an extended pH range in the 4-10 pH range when both buffer systems are combined, highlighting the tailoring of the method, influenced by buffer agents’ pKas.

#### pH-Dependent Growth Patterns of Recombinant *Chlamydomonas* Strains

The growth patterns of three recombinant *Chlamydomonas* strains— Cin, Cre, and Cpa—were analyzed on phosphate media (HSM nitrate) plates containing a stable pH gradient ranging from <4 acidic to >10 basic regions (**Figure 2**). **Figure 2, Panel A** shows the growth of each strain across the pH gradient. Each strain exhibited distinct pH tolerance profiles. Cin grew primarily in the slightly lower pH regions to somewhat neutral, indicating a preference for a mesophilic condition. In contrast, *C*re demonstrated tolerance to a broader pH range, with notable growth extending into neutral and slightly basic regions. Cpa displayed growth on a more alkaline pH range, including high pH regions, indicating a tolerance for alkaline conditions. **Figure 2, Panel B** demonstrates the competitive growth of Cin and Cre strains when mixed at a 1:1 ratio and evenly spread across the pH gradient. Boxed areas are where colonies could tolerate higher pH conditions, with magnified insets showing isolated colonies distributed in the transition zones. **Figure 2, Panel C** illustrates the competitive growth of the Cpa and Cre strains, mixed at a 1:1 ratio and spread across the pH gradient. Magnified insets provide closer views of the transition regions, showing isolated colonies distributed according to pH-dependent tolerance.

**Figure 2:**
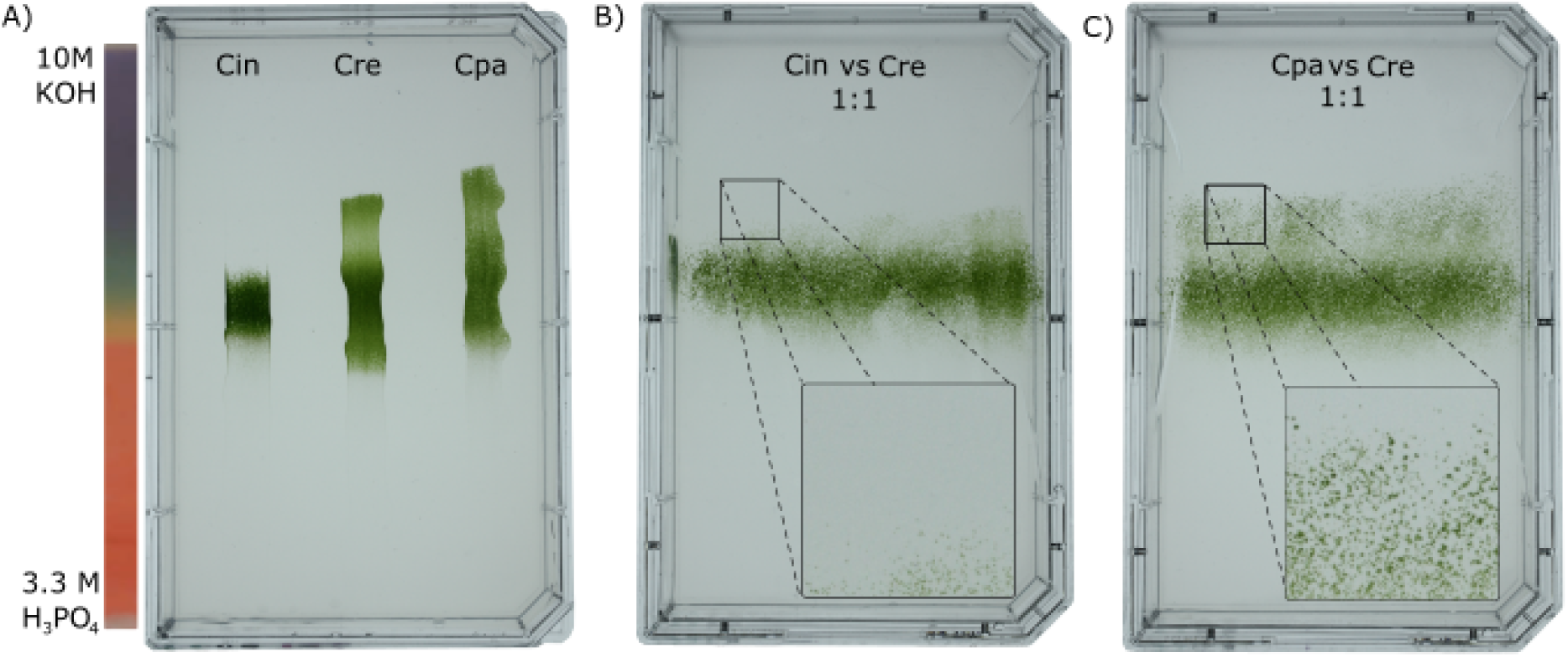
Growth patterns of recombinant *Chlamydomonas* strains on a pH gradient formed on phosphate media. **(A)** Growth of individual *Chlamydomonas* species on a pH gradient. Three recombinant strains—*Chlamydomonas incerta* (Cin), *Chlamydomonas reinhardtii* (Cre), and *Chlamydomonas pacific* (Cpa)—were applied to separate regions of the plate (100 µL patches) and allowed to grow. Cin harbors the vector pAH04mCherry, which confers bleomycin resistance; Cre harbors pJPCHx1_mVenus, which provides tolerance to hygromycin; and Cpa harbors pJP32PHL7, which also confers bleomycin resistance. The strains exhibited distinct pH tolerance profiles, as demonstrated by their spatial distribution along the acidic (red) to basic (blue) axis. **(B)** Competitive growth of Cin and Cre (1:1 mixture) across the pH gradient. A mixture of Cin and Cre cells (1:1 ratio) was uniformly spread across the entire plate. Solid boxes highlight the edge of tolerance to high pH for the mixture. Insets provide magnified views of these areas, showing isolated colonies by pH-dependent tolerance. **(C)** Competitive growth of Cpa and Cre (1:1 mixture) across the pH gradient. A mixture of Cpa and Cre cells (1:1 ratio) was uniformly spread across the plate. Solid boxes highlight the edge of tolerance to high pH. Magnified insets provide a closer view of the transition zones, showing isolated colonies by pH-dependent tolerance.

In addition to the observed growth patterns, colony recovery and selection experiments highlight *Chlamydomonas* strains’ competitive fitness in the gradient’s higher pH zones (**Supplementary Figure 3**). Panel A shows the number of colonies that recovered from the transition zone in Cin co-culture and Cre. Colony counts revealed a significantly higher recovery of Cre colonies than Cin colonies, indicating Cre’s superior survival and competitive ability in the higher pH transition zone. Panel B presents the co-culture results between Cpa and Cre. Here, Cpa demonstrated significantly higher colony recovery, particularly in the basic pH regions, reflecting its competitive advantage and tolerance for alkaline conditions. These findings confirm the pH-dependent survival and competitive interactions between strains, and showcase the ability of the method to distinguish among different strains based on the pH stress factor.

### Salinity Gradient plates

#### Method characterization with strains

The growth patterns of the three recombinant *Chlamydomonas* strains, Cre, Cin and Cpa were analyzed on a NaCl gradient plate created through replacing a section of the agar media with a concentrated NaCl (5M) agarose solution **(Figure 3**). A schematic representation of the experimental setup is shown in Panel A, where a section of agar was removed near one edge of the plate. Panel B illustrates the plate after agar removal, leaving a clean region for subsequent replacement with molten NaCl-rich agarose. Panel C shows the preparation of the NaCl gradient by replacing the removed section with 1.5% molten agar containing 5 M NaCl, which solidified and created a gradient as diffusion occurred. Panel D presents the growth of the three *Chlamydomonas* strains on the NaCl gradient. Distinct growth patterns were observed across the plate. Cin grew primarily in the low-NaCl regions, indicating sensitivity to high salt concentrations. Cre showed moderate tolerance to increased NaCl levels, with growth extending further into the gradient. In contrast, Cpa exhibited the greatest tolerance to high NaCl concentrations, with growth patches visible closer to the high-salt region of the plate. These results highlight the differential NaCl tolerance of the recombinant strains, with Cpa demonstrating superior adaptation to high-salt environments compared to Cre and Cin.

**Figure 3:**
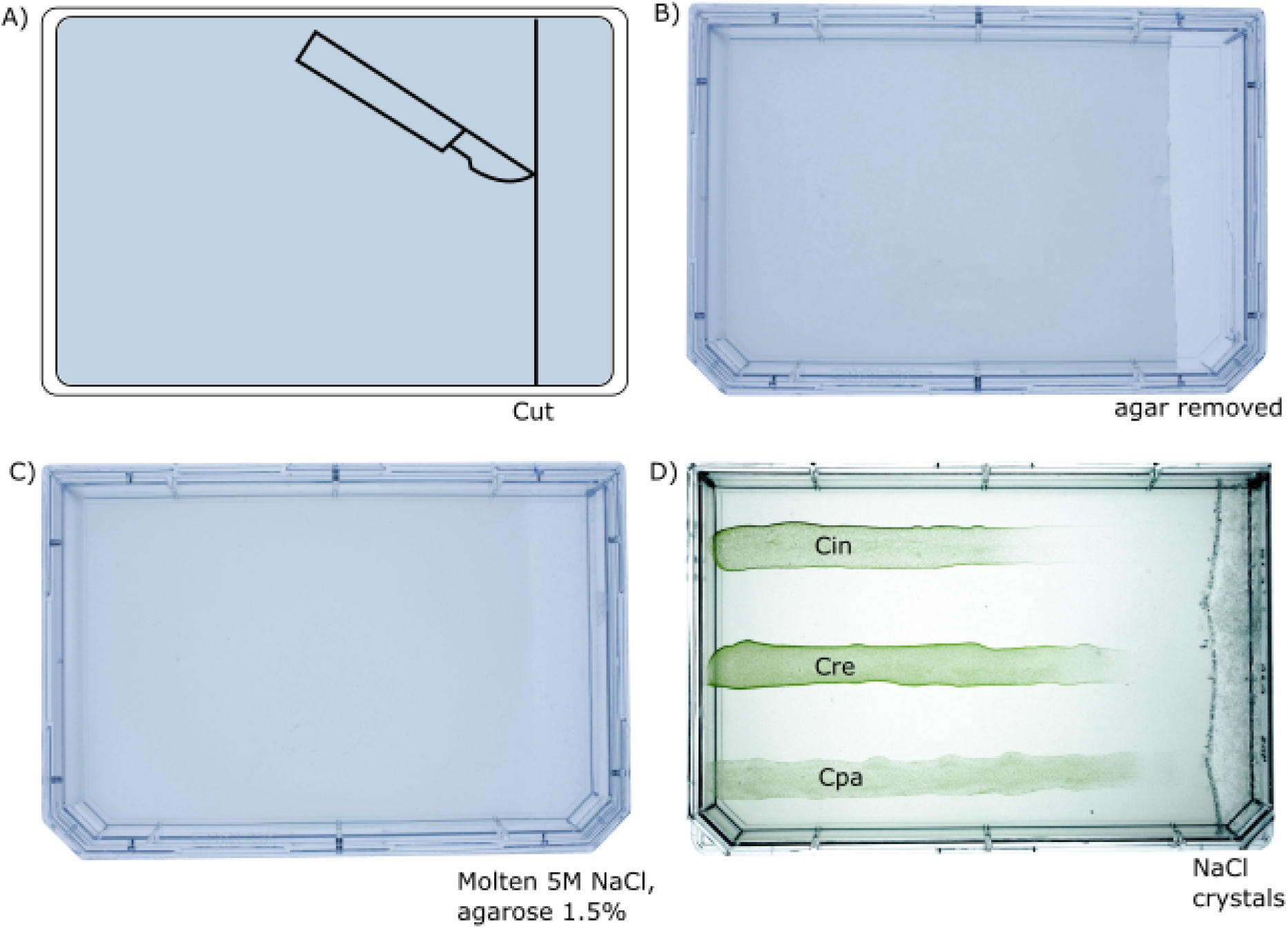
Growth of recombinant *Chlamydomonas* strains on a NaCl gradient plate prepared by media replacement. **(A)** Schematic representation of the experimental setup. A section of the agar media on the plate was removed by making a straight cut with a scalpel near one edge of the plate. **(B)** Image of the plate after the agar was removed, leaving a clean region for subsequent replacement with NaCl-rich agar. **(C)** Preparation of the NaCl gradient. The removed section was replaced with molten 1.5% agarose containing 5 M NaCl, which was allowed to solidify, creating a NaCl gradient across the plate as diffusion occurred. **(D)** Growth patterns of the recombinant *Chlamydomonas* strains—*Chlamydomonas incerta* (Cin), *Chlamydomonas reinhardtii* (Cre), and *Chlamydomonas pacific* (Cpa)—on the NaCl gradient. Each strain was inoculated in 100 µL patches near the low-NaCl region of the plate. Cin harbors the vector pAH04mCherry, which confers resistance to bleomycin; Cre harbors pJPCHx1_mVenus, which confers resistance to hygromycin; and Cpa harbors pJP32PHL7, which confers resistance to bleomycin.

To validate the findings in the gradient plate assay, normalized chlorophyll fluorescence was measured as a proxy for growth under varying NaCl concentrations in liquid media (**Figure 4**). The strains were cultured in TAP media supplemented with increasing concentrations of NaCl (0–400 mM) and incubated for 7 days under continuous light with agitation. Cin exhibited a sharp decline in fluorescence beyond 100 mM NaCl, indicating sensitivity to elevated salt concentrations. Cre demonstrated moderate tolerance, maintaining measurable fluorescence up to 200 mM NaCl before declining. In contrast, Cpa displayed the highest salt tolerance, retaining substantial fluorescence even at 300 mM NaCl, though it declined at 400 mM. These results corroborate the growth patterns observed on the NaCl gradient plates, further confirming Cpa as the most salt-tolerant strain, followed by Cre, with Cin exhibiting the lowest tolerance to NaCl stress.

**Figure 4:**
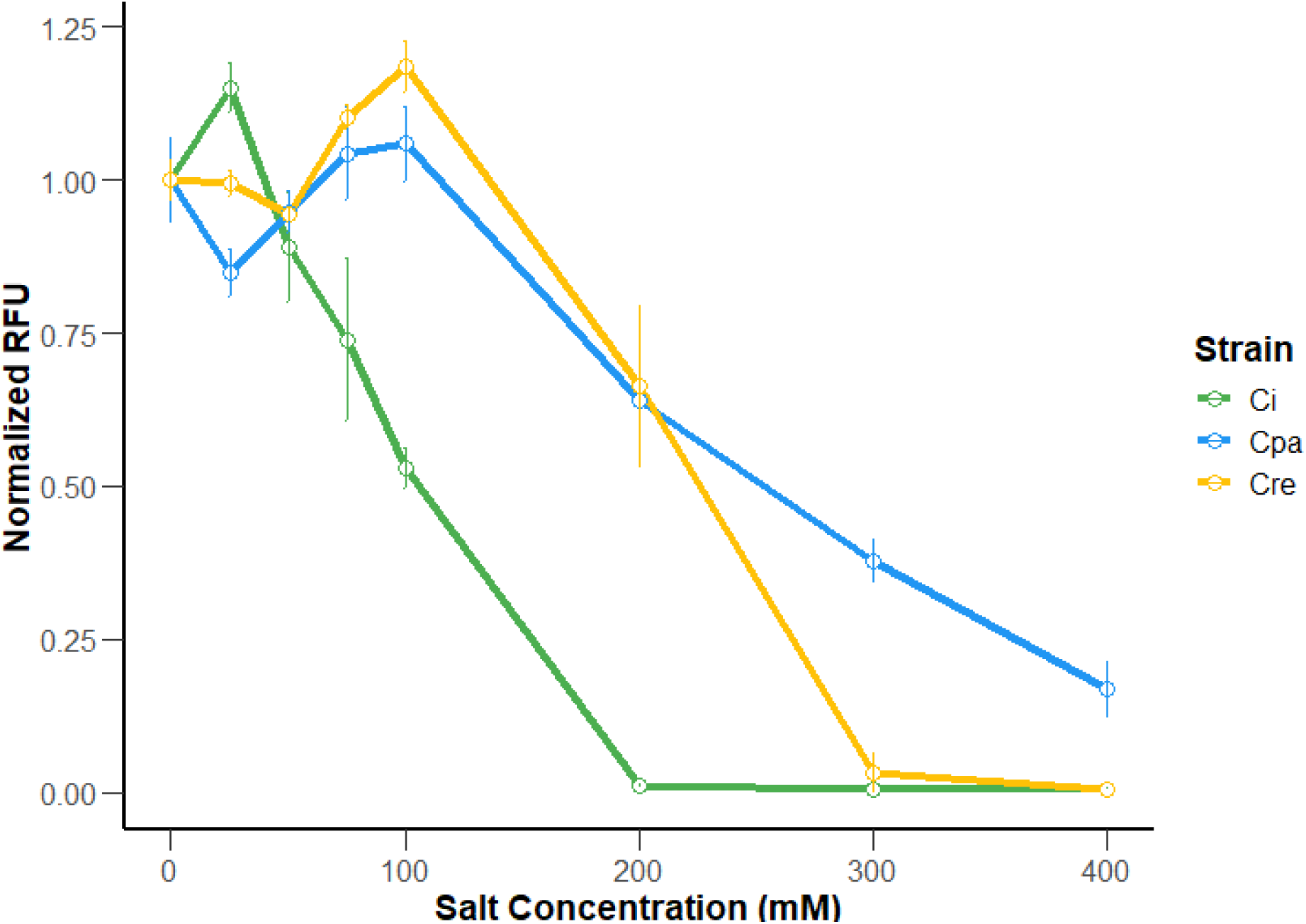
Normalized chlorophyll fluorescence of recombinant *Chlamydomonas* strains under varying NaCl concentrations. Normalized relative fluorescence units (RFU) were measured for three recombinant *Chlamydomonas* strains: *Chlamydomonas incerta* (Cin), *Chlamydomonas pacifica* (Cpa), and *Chlamydomonas reinhardtii* (Cre). Cin harbors the vector pAH04mCherry (conferring bleomycin resistance), Cpa harbors pJP32PHL7 (conferring bleomycin resistance), and Cre harbors pJPCHx1_mVenus (conferring hygromycin resistance). Cells were inoculated into a 96-well plate containing TAP media supplemented with NaCl at increasing concentrations (0–400 mM). The cultures were incubated under continuous light and agitation for 7 days in four replicas per condition. The graph depictst the mean of four replicas, and the error bars the standard deviation. Chlorophyll fluorescence, as a proxy for growth, was measured and normalized to the initial fluorescence value in TAP media without additional NaCl (0 mM).

#### Competitive tolerance assay of *Chlamydomonas* strains on NaCl gradient plates

The competitive growth of recombinant *Chlamydomonas* strains was analyzed on NaCl gradient plates under co-culture conditions to assess their tolerance to varying salt concentrations and validate the method to distinguish among strains (**Figure 5**). **Figure 5, Panel A** shows the growth of a 1:1 mixture of Cin and Cre. The strains were evenly spread across the NaCl gradient plate, and growth patterns were examined under competitive conditions. **Figure 5, Panel B** presents the competitive growth of Cpa and Cre in a 1:1 mixture on a NaCl gradient plate. The magnified inset highlights the edge of the tolerance zone for both panels. The edge of the tolerance zone was scraped to perform the colony recovery experiment. Colonies from the competitive co-culture of Cin and Cre were recovered, resuspended, and plated under selective conditions to sort between Cre Hyg^R^ or Cin Zeo^R^ colonies (**See Method section**). Colony counts revealed a significantly higher (*p-value < 0.001*, ANOVA) recovery of Cre colonies compared to Cin, confirming Cre’s enhanced survival in the NaCl gradient (**Supplementary Figure 4**). Similarly, in the competitive co-culture of Cpa Zeo^R^ and Cre Hyg^R^, colony recovery demonstrated a significantly higher abundance of Cpa colonies, reflecting its superior fitness and tolerance to higher NaCl levels. These results collectively show that *Cre* exhibits greater salt tolerance than *Cin* while *Cpa* demonstrates the highest tolerance to elevated NaCl concentrations.

**Figure 5:**
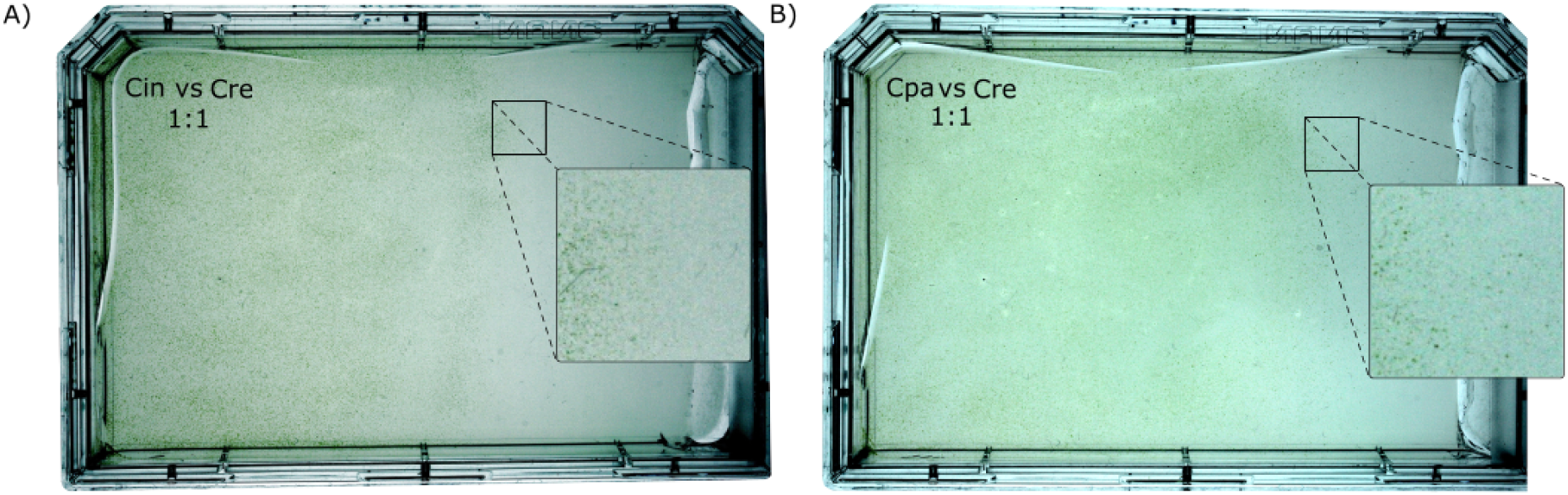
Competitive growth of recombinant *Chlamydomonas* strain mixtures on NaCl gradient plates. **(A)** Growth of a 1:1 mixture of *Chlamydomonas incerta* (Cin) and *Chlamydomonas reinhardtii* (Cre) on a NaCl gradient plate. Cin harbors the vector pAH04mCherry (conferring bleomycin resistance), while Cre harbors pJPCHx1_mVenus (conferring hygromycin resistance). The strains were evenly spread across the plate near the low-NaCl region and allowed to grow under competitive conditions. The magnified inset highlights the edge of the tolerance zone. **(B)** Growth of a 1:1 mixture of *Chlamydomonas pacific* (Cpa) and *Chlamydomonas reinhardtii* (Cre) on a NaCl gradient plate. Cpa harbors the vector pJP32PHL7 (conferring bleomycin resistance), while Cre harbors pJPCHx1_mVenus (conferring hygromycin resistance). The strains were evenly spread near the low-NaCl region. The magnified inset highlights the edge of the tolerance zone.

### Temperature Tolerance assay

#### Temperature gradient using a thermocycler

The temperature tolerance of the *Chlamydomonas* strains was assessed using a temperature gradient assay (**Figure 6**). Cultures of each strain were dispensed into a PCR plate containing TAP media and subjected to a temperature gradient ranging from low (34°C) to high (42°C). The plates were incubated for 24 hours to evaluate tolerance under thermal stress conditions. Following incubation, the cultures were resuspended and spotted onto TAP media agar plates, and growth was monitored over 7 days to determine strain-specific temperature tolerance profiles.

**Figure 6:**
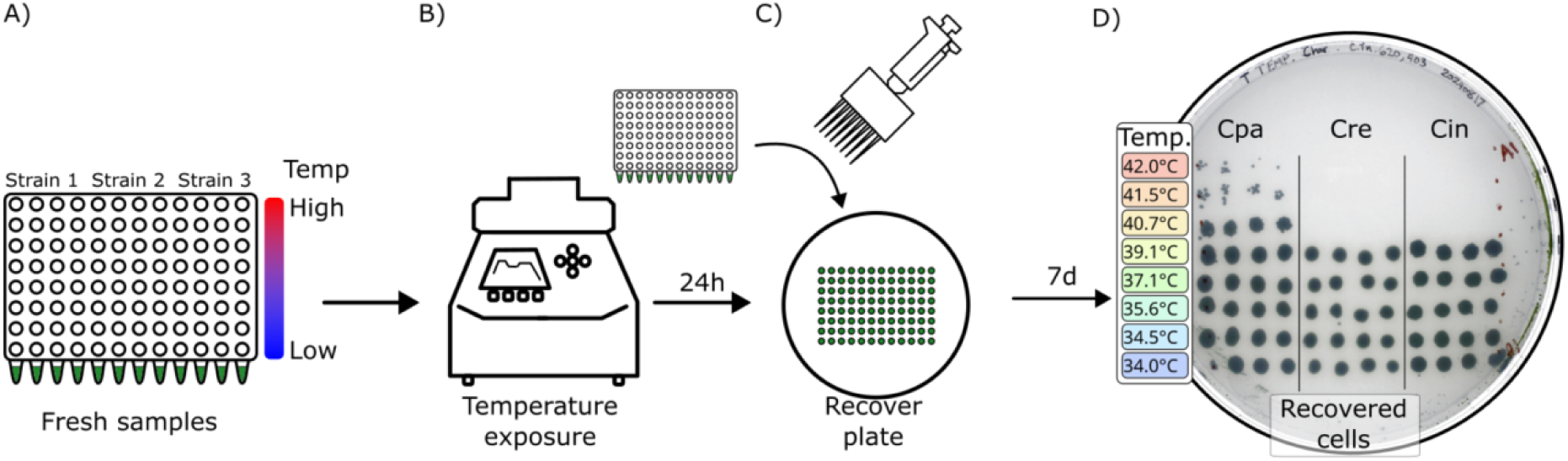
Temperature-tolerance assay of recombinant *Chlamydomonas* strains. **(A)** Experimental setup for assessing temperature tolerance. Cultures of *Chlamydomonas pacific* (Cpa), *Chlamydomonas reinhardtii* (Cre), and *Chlamydomonas incerta* (Cin) were prepared in TAP media and dispensed into a PCR plate (100 µL per well), with four replicates per temperature condition. A gradient of temperatures, ranging from low (blue) to high (red), was applied across the rows of the plate. **(B)** The PCR plate was sealed with a transparent film and incubated in a thermocycler for 24 hours to check cells tolerance at the designated temperature gradient. **(C)** Following incubation, cultures were resuspended using a multichannel pipette, and 5 µL from each well was spotted onto a TAP media agar plate to assess growth under uniform conditions. **(D)** Growth of the three recombinant strains after 7 days of incubation. The spotted samples reveal distinct temperature tolerance profiles for each strain. Cpa (left section), Cre (middle section), and Cin (right section) display varying growth intensities and viability across the temperature gradient, reflecting their specific thermal adaptation ranges.

Distinct growth patterns emerged across the temperature gradient. Cpa exhibited the highest thermal tolerance, showing robust tolerance at elevated temperatures. In contrast, Cre and Cin displayed moderate thermal tolerance, with growth restricted to the mesophilic temperature range and reduced viability at the highest temperatures. These findings highlight significant differences in temperature adaptation among the strains, with Cpa demonstrating superior thermal resilience compared to Cre and Cin.

#### Competitive Tolerance Assay of *Chlamydomonas* Strains After Temperature Screening

To evaluate the capacity of the method to distinguish different strains, a competitive tolerance assay for the *Chlamydomonas* strains was performed with the strain mixtures, then spotted onto non-selective and selective TAP media (**Figure 7**). The selective plates allowed us to distinguish between the mixed strains. The left plate (TAP Ctrl) shows growth on non-selective media, where mixtures of Cpa and Cre at both 1:1 and 1:100 ratios, as well as Cin and Cre at a 1:1 ratio, were tested at different temperatures. Growth of all strains was visible under these conditions, indicating survival of at least one of the strains following temperature exposure. On the middle plate (TAP Zeo), which was supplemented with 15 µg/mL zeocin to select for bleomycin-resistant strains (Cpa and Cin), growth was observed at the temperatures expected to each respective strain. Notably, Cpa colonies were detectable even at the 1:100 ratio in competition with Cre, highlighting the method’s capacity to select low-abundant strains in a sample. The right plate (TAP Hyg), supplemented with 30 µg/mL hygromycin to select for hygromycin-resistant strains (Cre), showed growth exclusively for Cre in the expected temperatures (< 39.1°C). These results demonstrate the competitive fitness of *Chlamydomonas* strains with Cpa displaying strong tolerance to temperature, and the method was capable of selecting Cpa strains, even at low proportions, based on temperature.

**Figure 7:**
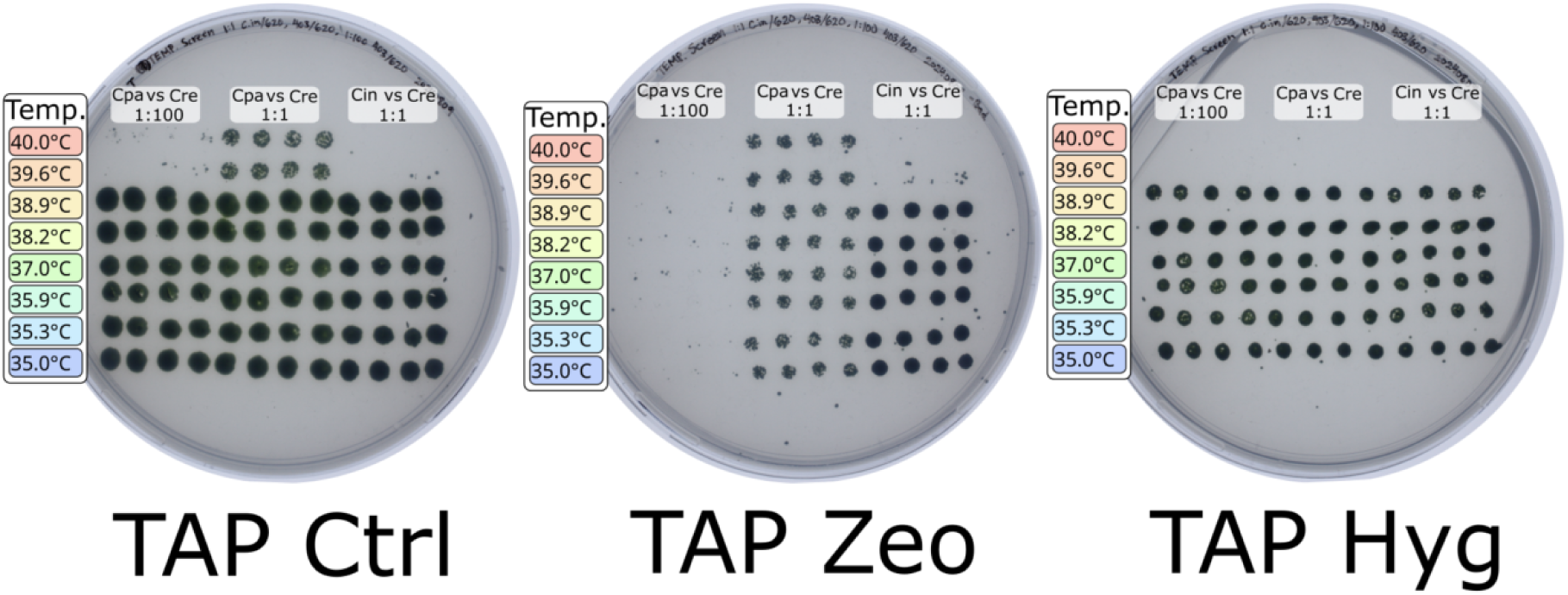
Competitive growth of *Chlamydomonas* strain mixtures after temperature screening, plated on selective and non-selective media. Left Plate (TAP Ctrl): Cells from the temperature screening assay, including mixtures of *Chlamydomonas pacific* (Cpa) and *Chlamydomonas reinhardtii* (Cre) at 1:1 and 1:100 ratios, as well as *Chlamydomonas incerta* (Cin) and Cre at 1:1 ratio, were spotted on non-selective TAP media. Each mixture was plated in 5 µL drops. **Middle Plate (TAP Zeo):** The same cell suspensions were plated on TAP media supplemented with 15 µg/mL zeocin, selecting for Cpa and Cin (bleomycin-resistant strains). In the 1:1 mixtures of Cpa vs Cre and Cin vs Cre, growth is visible only for Cpa and Cin. In the 1:100 mixture of Cpa and Cre, Cpa colonies were detectable even at this low proportion, demonstrating the ability to select for Cpa in high temperatures even in lower proportions. **Right Plate (TAP Hyg):** Cells were plated on TAP media supplemented with 30 µg/mL hygromycin, selecting for Cre (hygromycin-resistant strain). In the 1:1 mixtures of Cpa vs Cre and Cin vs Cre, growth is visible only for Cre. Similarly, in the 1:100 mixture of Cpa and Cre, only Cre colonies were observed at the limit temperature tolerance for the strain.

### Light Tolerance Assay

#### Characterization of the *Chlamydomonas* Strains

To demonstrate the efficacy of a light tolerance assay in distinguishing algae strains, we show its application on the three *Chlamydomonas* strains exposed to a gradient of light intensities (**Figure 8**). A schematic representation of the experimental setup is shown in Panel A, where two TAP media plates were prepared for light exposure. One plate was partially shaded using a paper filter, creating a variation ranging from ∼40 to ∼3000 µE/m²s (shaded plate). In contrast, the second plate was exposed directly to high-intensity LED light, generating a gradient from ∼1200 to ∼3000 µE/m²s (direct light plate), with a control region at 500 µE/m²s. Panels B and C display the growth patterns of the strains under shaded and direct light conditions. On the shaded plate, robust growth was observed for all three strains across the lower light intensities (∼40–500 µE/m²s). However, significant differences emerged among the strains on the direct light plate. Cpa exhibited the highest light tolerance, with tolerance at >1200 µE/m²s. Cre displayed low tolerance, with colonies emerging in a short region near the 500 µE/m²s control region but absent at the highest intensities. In contrast, Cin demonstrated moderate tolerance, with colonies emerging in regions ∼1200 µE/m²s. These results confirm that the light tolerance assay effectively distinguishes *Chlamydomonas* strains based on their ability to tolerate varying light intensities, allowing for the identification of strain-specific light tolerance capabilities.

**Figure 8:**
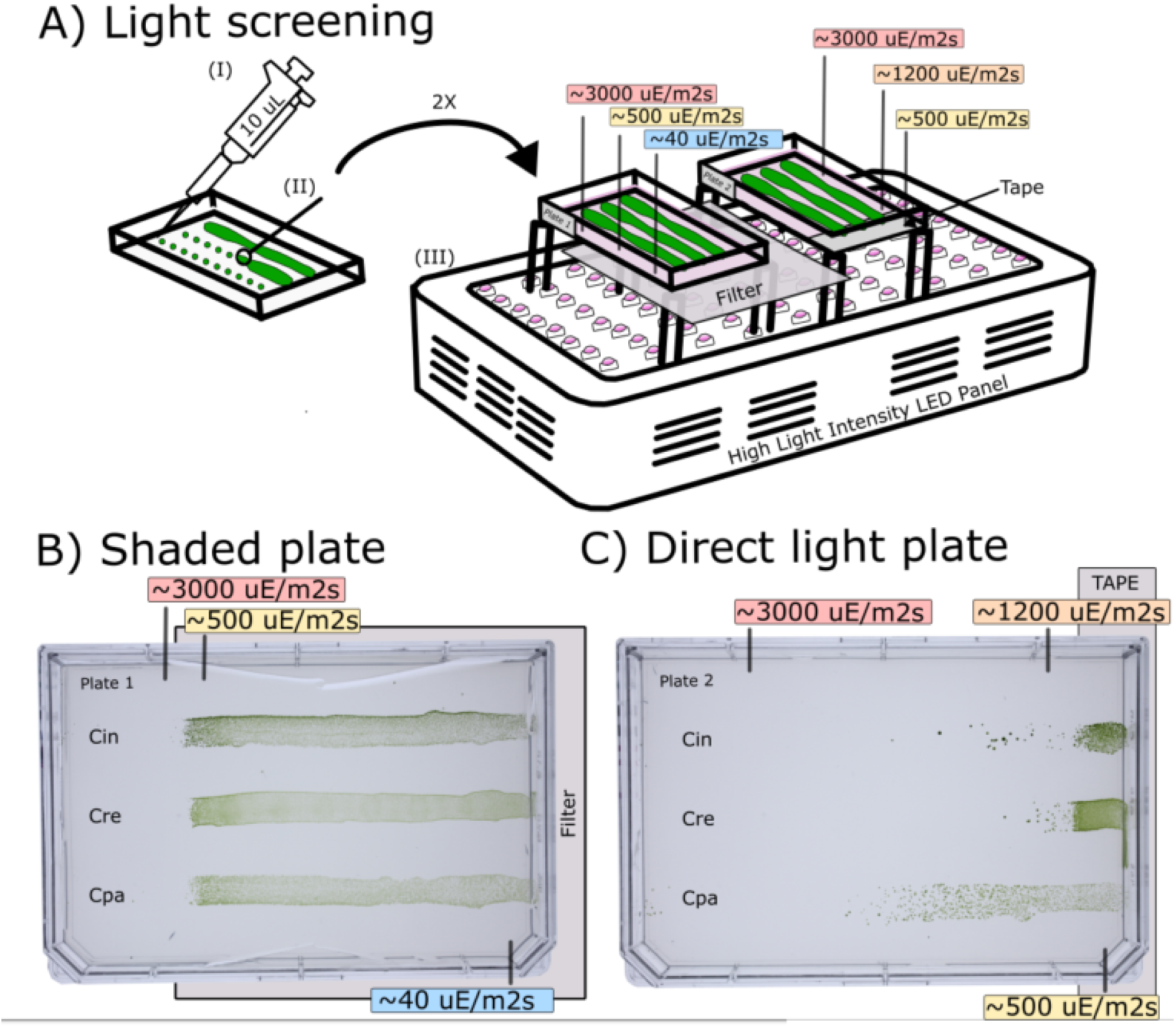
Light tolerance assay of recombinant *Chlamydomonas* strains under gradient light exposure. **(A)** Schematic representation of the experimental setup. Recombinant strains *Chlamydomonas incerta* (Cin), *Chlamydomonas reinhardtii* (Cre), and *Chlamydomonas pacific* (Cpa) were patched onto TAP media plates (100 µL per strain, side by side) in linear streaks. Cin harbors the vector pAH04mCherry, which confers resistance to bleomycin; Cre harbors pJPCHx1_mVenus, which confers resistance to hygromycin; and Cpa harbors pJP32PHL7, which also confers resistance to bleomycin. Two plates were prepared for the light exposure assay. The first plate was partially shaded using a paper light filter, creating a gradient of light intensities ranging from ∼40 µE/m²/s in the shaded area to ∼3000 µE/m²/s in the exposed region. The second plate was exposed directly to a high-intensity LED panel without a filter, generating a gradient of light intensities from ∼500 µE/m²/s to ∼3000 µE/m²/s. A portion of each plate was covered with tape to act as a control, ensuring the inoculated cells remained viable after light exposure. Cells were exposed to the light gradient for 24 hours, followed by 7 days of recovery under standard growth conditions. **(B)** Shaded plate: Growth patterns of Cin, Cre, and Cpa under a gradient of light intensities created by the paper filter. Robust growth was observed in the shaded regions (∼100–500 µE/m²/s), while exposure to the highest intensity region (∼3000 µE/m²/s) resulted in significant growth inhibition or bleaching for all three strains. **(C)** Direct light plate: Growth patterns of Cin, Cre, and Cpa under a direct, unfiltered light gradient. The strains exhibited differential tolerance to light intensities, with inhibition observed at the highest intensity (∼3000 µE/m²/s). Moderate growth was observed in the intermediate intensity zones (∼500–1200 µE/m²/s). The tape-protected region on both plates confirmed that all inoculated cells were viable under standard growth conditions.

#### Screening and selection of *Chlamydomonas* strains using the light tolerance method

The light tolerance assay was used to screen mixed cultures of the *Chlamydomonas* strains under a light intensity gradient to identify strains capable of surviving high-light conditions (**Figures 9 and Supplementary Figure 5**). Mixtures of Cpa and Cre, as well as Cin and Cre, were co-inoculated at a 1:1 ratio and exposed to increasing light intensities ranging from ∼500 to ∼3000 µE/m²/s for 24h. Colonies that survived in the high-light regions, particularly within the transition zones, were isolated, grown in 96-well plates, and subsequently dotted onto selective and non-selective media for strain identification. We used the non-selective TAP media (TAP Ctrl) to infer the number of successfully picked colonies in our assay. On TAP Zeo, supplemented with 15 µg/mL zeocin to select for bleomycin-resistant strains, colonies of Cpa and Cin were recovered, while Cre colonies were absent. Conversely, on TAP Hyg, supplemented with 30 µg/mL hygromycin to select for Cre (hygromycin-resistant strain), only Cre colonies were observed, with Cpa and Cin colonies eliminated. These colonies were picked into 96-well plates, and after 7 days of growth, the cultures in each well were stamped onto a non-selective TAP agar plate, a TAP Hyg plate, or a TAP Zeo plate. After 7 days of growth, these plates revealed that in the Cpa and Cre mixture, only 3 out of 68 recovered colonies were Cre, while the remaining 65 were Cpa. Similarly, in the Cin and Cre mixture, only 3 out of 85 colonies were Cre, with the remaining 82 Cin. Starting from an initial ratio of 1:1 (Cre:Cpa and Cre:Cin), the post-selection frequencies showed a depletion of Cre colonies (Cpa mix: 3 out of 68 colonies, Exact binomial test, p < 0.0001; Cin mix: 3 out of 85 colonies, Exact binomial test, p < 0.0001). These results demonstrate that Cpa and Cin strains exhibit superior survival in high-light intensity zones compared to Cre, as observed in **Figure 8**. The light tolerance assay effectively distinguishes strains within mixed cultures, underscoring its reliability for identifying and isolating light-tolerant strains.

**Figure 9:**
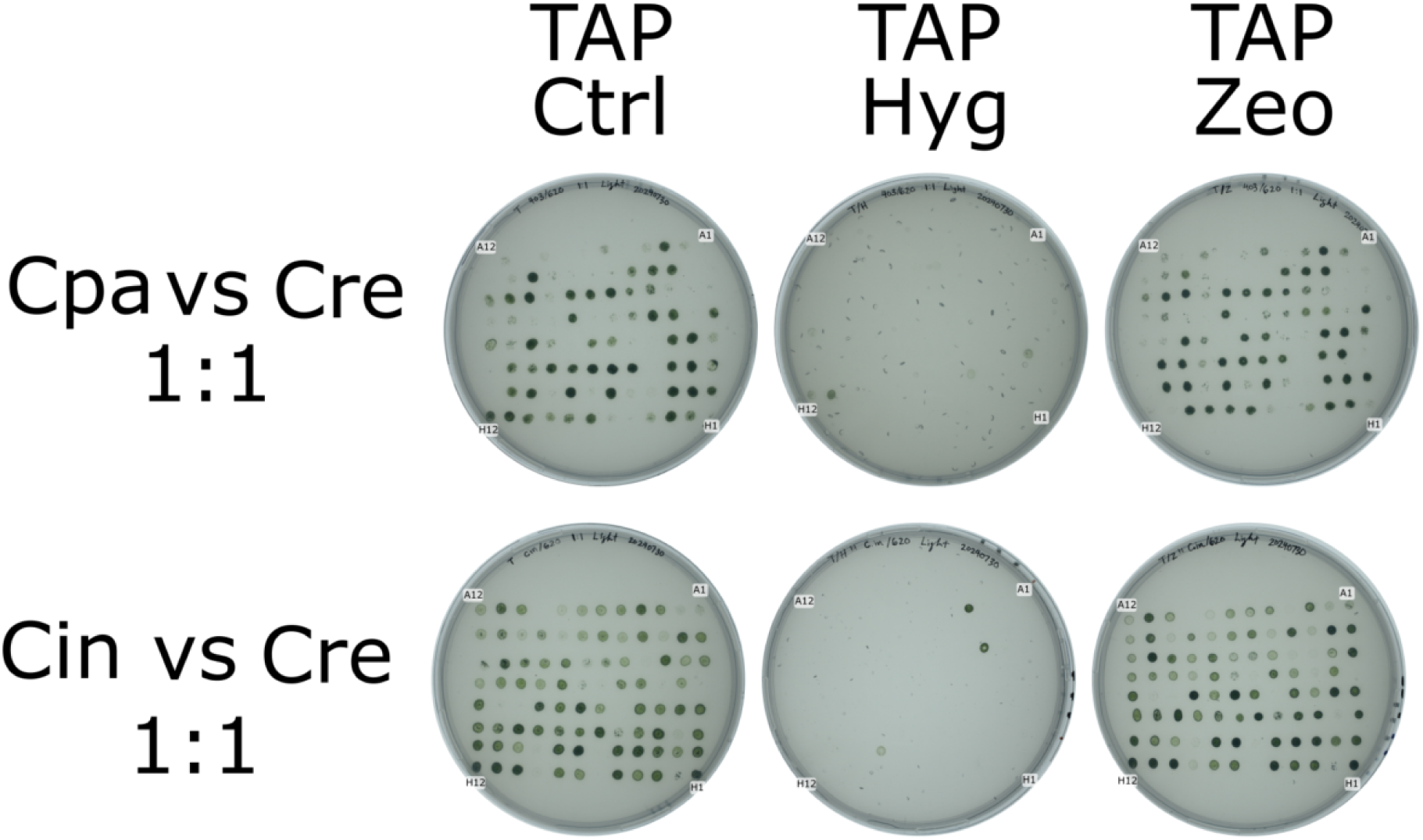
Selection of *Chlamydomonas* strains from colonies grown in high-light regions under competitive conditions. Colonies that survived in the higher-light intensity region of the light gradient plates were picked and grown in a 96-well plate in TAP liquid media. Subsequently, 5 µL of each well was spotted onto TAP agar plates with different selection conditions: non-selective TAP media (TAP Ctrl), TAP with 15 µg/mL zeocin (TAP Zeo), and TAP with 30 µg/mL hygromycin (TAP Hyg). **Top Row (Cpa vs Cre 1:1):** Results for colonies derived from the 1:1 mixture of *Chlamydomonas pacific* (Cpa) and *Chlamydomonas reinhardtii* (Cre). On TAP Zeo, colonies corresponding to the bleomycin-resistant strain Cpa (pJP32PHL7) were recovered, while colonies corresponding to Cre (hygromycin-resistant, pJPCHx1_mVenus) were eliminated. Conversely, colonies of Cre were selectively recovered on TAP Hyg. **Bottom Row (Cin vs Cre 1:1):** Results for colonies derived from the 1:1 mixture of *Chlamydomonas incerta* (Cin) and *Chlamydomonas reinhardtii* (Cre). Similar to the Cpa vs Cre competition, growth was observed on the TAP Ctrl plate for most colonies. Cin (bleomycin-resistant, pAH04mCherry) colonies were selectively recovered on TAP Zeo, while Cre colonies were eliminated. On TAP Hyg, Cre colonies dominated, confirming the presence of Cre in the high-light region.

## Discussion

In this study, we developed and validated a set of low-cost screening methods capable of evaluating the tolerance of algae strains to extreme environmental conditions, including pH, salinity, temperature, and light. These agar plate-based techniques, combined with simple diffusion gradients and selective recovery assays, provide accessible tools for distinguishing strains with desirable stress-tolerance traits. By demonstrating the effectiveness of these methods in identifying strain-specific responses and competitive fitness under varying conditions, we highlight their utility in streamlining the bioprospecting and characterization of algae. Importantly, these approaches are adaptable to various microorganisms beyond algae, enabling broader applications in biotechnology, microbial ecology, and strain selection for industrial purposes. The ability to screen for traits such as pH, salt, temperature, and light tolerance at a reduced cost and technical requirement makes these methods particularly valuable for laboratories with limited resources, fostering advancements in algae-based biotechnologies globally. Aside from bioprospecting, these methods have the potential to be combined with breeding programs to enhance traits in already industry-relevant strains and improve their robustness. In fact, we exploited *Cpa* mating capacity coupled with mutagenesis to further improve its light tolerance (18), demonstrating the applicability of the methods described here.

Furthermore, phenotypic screening is greatly enhanced with modern genomic and genetic tools. Identifying a stress-tolerant strain is just the first step; understanding *why* it is tolerant opens the door to rational improvement and predictive breeding. Fortunately, the decreasing cost of genome sequencing and advances in algal genetics now make it feasible to map the genetic basis of complex traits like stress tolerance. By combining our high-throughput screening assays with genome-wide analyses, we can accelerate trait discovery in microalgae. One approach is to perform genome-wide association studies (GWAS) on a mutant pool selected for each specific trait and compare it with the parents. By genotyping or sequencing these strains, statistical associations can be made between genetic markers and the tolerance trait phenotype to identify Quantitative Trait Loci (QTL): genomic regions linked to the trait (19). This strategy, widely used in crop plants, can pinpoint specific genes or mutations that confer improved stress resistance. Those genes become targets for further validation (through gene expression studies or knock-out/overexpression in model algae) and marker-assisted strain breeding. The QTL approach allows one to dissect the genetic and physiological components underlying abiotic stress tolerance, even for polygenic traits (19). In algae, where classical breeding is less established, GWAS on natural or laboratory-evolved variants could rapidly yield candidates for the molecular basis of heat, light, pH, or salinity tolerance, with implications for crop plants, as homologous gene candidates can quickly be found.

### pH Tolerance

Ambient pH affects microalgal growth in multiple direct and indirect ways. The hydrogen-ion concentration of the medium governs the solubility and availability of essential nutrients (such as inorganic carbon and metals). It can therefore limit uptake of CO₂ or minerals if not in the optimal range (20). Extremely low or high pH also has direct cytoplasmic effects: it can lead to gross alterations in membrane transport processes and metabolic functions as cells struggle to maintain internal pH homeostasis (21), as effects on secreted and exposed proteins. Most microalgae grow best in near-neutral pH conditions; significant deviations can inhibit enzyme activities, destabilize membrane potential, and cause nutrient precipitation or toxicity.

Assessing pH tolerance in algae is complicated by the fact that microalgal metabolism itself can alter the medium pH. For example, active CO₂ fixation or ammonia assimilation raises the culture pH. At the same time, respiration or nitrate uptake can lower it, meaning a culture’s pH may drift over time despite initial adjustment (22). Many traditional pH tolerance assays only capture short-term survival at a given pH and may not reflect an alga’s ability to acclimate to prolonged pH stress (23). Additionally, maintaining stable pH in experiments is non-trivial; buffer systems can be overwhelmed by biological activity, and expensive apparatus can be required to keep one pH target (24). This underscores the need for simple and reliable methods, such as gradient approaches, to reliably evaluate and select for pH-stress-tolerant strains.

We successfully established a robust, cost-effective, and versatile screening method to evaluate algae strain tolerance under different pHs in a simple and low-cost manner. This method utilized simple diffusion-based pH gradients on agar plates, inspired by previous methods (25,26). In our assay, the stability of the gradients was maintained over 14 days, enabling a comprehensive evaluation of pH tolerance and competitive fitness in individual and co-culture settings. The use of a single pH change inducer creates a dynamic variation of pH across the plate (**Supplementary Video 2**). This approach effectively identified pH-specific growth patterns and highlighted strain-specific competitive advantages, demonstrating its potential utility for bioprospecting and strain improvement efforts. Our system provides a stable, well-defined gradient supported by time-lapse analysis with pH indicators **(Figure 1**), live-cell testing (**Figure 2**), and competitive fitness validation (**Supplementary Figure 3**). These features uniquely position this method as an innovative tool for identifying resilient strains within mixtures, enabling high-throughput screening of stress-tolerant algae and other microorganisms.

However, during early development, we encountered an unexpected challenge when ammonium-based media (NH₄⁺) were used, as the formation of ammonia gas (NH₃) caused pH disruptions, nitrogen availability disruption, and potential toxicity. Replacing ammonium with nitrate from the media mitigated this issue, highlighting the importance of media composition consideration in experimental setups and industrial settings.

Additionally, our results demonstrated the method’s ability to distinguish between two strains when present in equal proportions. However, an intriguing and unexplored strategy is the sequential screening of mixed pools containing strains at uneven abundances, using an enrichment strategy by applying selection iteratively. This approach could be beneficial for highly tolerant strains, potentially enhancing the sensitivity and resolution of the method and enabling the identification and isolation of strains present at initially low abundances.

Furthermore, the current study was limited to three algae strains tested in a simple media system, which may not represent the full range of potential physiological responses. For species with complex nutrient requirements or sensitivity to pH shifts, additional factors, such as NH₄⁺ toxicity, could influence results. The method is also constrained to algae that can grow on solid media, potentially excluding strains with specific growth requirements.

To further expand the utility of this method, future research should explore a broader range of buffer combinations to fine-tune pH gradient dynamics. Incorporating advanced imaging techniques could improve gradient resolution and visualization. Building on these results, this method could be employed to develop algae strains with enhanced pH tolerance through directed evolution or breeding programs, ultimately contributing to advancements in algae-based biotechnology.

### Salinity Tolerance

The ability to maintain and adapt to optimal salinity conditions is crucial for the survival and growth of algae, as salinity directly impacts osmotic balance, ion homeostasis, and cellular metabolism. Different species exhibit varying levels of tolerance and fitness to specific salinity ranges, reflecting their ecological origins and adaptability to freshwater, brackish, or marine environments. When external salt concentrations rise, water tends to exit the cell, causing dehydration, and excessive Na⁺/Cl⁻ or other ions can disrupt cellular enzymes and membranes (ionic imbalance) (27). To cope with high salinity, microalgae activate osmoregulatory mechanisms: they accumulate compatible solutes (osmoprotectants) like sugars, polyols, amino acids, or quaternary ammonium compounds, and adjust internal ion concentrations to retain water and protect proteins (28).

Our results demonstrated that it is possible to separate and compare algae strains based on salinity tolerance using a simple agar-based method that creates a NaCl gradient through diffusion (**Figure 3 and 5**). Distinct growth patterns were observed among the three recombinant *Chlamydomonas* strains on NaCl gradient plates. Cpa exhibited the highest salt tolerance, thriving in the plate regions with elevated NaCl concentrations, followed by Cre. In contrast, Cin showed limited growth and was restricted to low-NaCl areas, highlighting its sensitivity to salt stress. These findings were corroborated by chlorophyll fluorescence assays in liquid media (**Figure 4**), further validating the utility of this gradient-based method for distinguishing salinity tolerance among algae strains.

The method offers significant advantages as a simple and effective screening tool for salinity tolerance in algae. Unlike traditional approaches, this technique employs a straightforward NaCl gradient by replacing a portion of agar with NaCl-rich agarose. The observed sensitivity of strains to the gradient is primarily driven by the inability of salt-intolerant cells to grow in high-salinity regions, where the NaCl concentration is inhibitory due to osmotic stress and ionic imbalances.

One challenge encountered was the formation of salt crystals on the plate due to evaporation, an issue related to the near-saturation levels of the 5M NaCl agarose solution. While this did not significantly impact the overall gradient stability or the ability to discern between strains, it highlights the physical limitations of using high salt concentrations and improving or discerning among highly extremophile strains such as *Dunaliella salina*, which naturally occurs in environments with >5.5 M salt content (29).

Furthermore, unlike the stable pH gradient system, the NaCl gradient is subject to continuous diffusion over time, which may reduce its effectiveness for slower-growing strains. Additionally, the method also requires strains capable of growing on solid media, which may exclude some species.

Future research should provide a more detailed characterization of the diffusion dynamics of NaCl within the agar system. This could involve employing conductivity measurements to monitor the diffusion dynamics and progression of the gradient. Although the formation of the salt gradient will inevitably follow Fick’s First Law of Diffusion and therefore be exploitable even on unknown salt diffusion dynamics, precisely quantifying these dynamics will enhance the reliability and reproducibility of the assay.

### Temperature Tolerance

Temperature strongly influences microalgal physiology by affecting biochemical reaction rates and the integrity of cellular structures. Like most organisms, microalgae have an optimum temperature range for growth; outside this range, enzymatic kinetics are suboptimal, and extreme temperatures can cause cellular damage. Elevated heat accelerates metabolic reactions up to a point, but beyond the optimum, it leads to protein misfolding/denaturation and increases membrane fluidity to potentially destabilizing levels (30,31). Key processes like DNA replication and photosynthesis become impaired – for instance, the D1 protein of Photosystem II is prone to heat-induced inactivation, and the overall photosynthetic rate drops as temperature rises above a strain’s comfort zone (31). In practical outdoor cultivation, temperature fluctuations can be a significant challenge. Shallow algal ponds or surface-exposed photobioreactors often heat up with daytime sun and cool off at night. It is not uncommon for culture temperatures to swing from about 20 °C in the early morning to 35–40 °C or more in the afternoon in warm climates (32). Such swings can induce daily cycles of stress: microalgae might experience temporary heat stress at noon (risking partial photoinhibition or growth arrest) and then suboptimal cool conditions at night. If a strain cannot tolerate these fluctuations – for example, if it irreversibly loses viability above 30 °C – the culture will fail under outdoor conditions. Selecting for thermotolerant strains is thus crucial.

Our findings demonstrate the utility of a thermocycler-based approach for assessing temperature tolerance in algae strains. Using this method, we identified distinct thermal tolerance profiles among Cpa, Cre, and Cin, with Cpa exhibiting the highest tolerance to elevated temperatures (**Figure 6**). The assay also effectively highlighted strain-specific survival in competitive co-culture conditions, demonstrating the robustness of the approach for distinguishing and selecting strains based on thermal resilience (**Figure 7**). The thermocycler assay operates by placing cells in PCR plate wells and exposing them to a controlled temperature gradient. After exposure, the cells are transferred to fresh media to determine survival and identify the critical temperature thresholds where cell death occurs (**Figure6**). This approach offers a simple yet effective mechanism for high-resolution analysis of thermal tolerance and screening.

During the study, the cells were kept in the dark during the thermocycler assay, which may reduce viability for some strains. Nonetheless, most algae can tolerate temporary darkness, as shown by the survival of the tested strains after incubation (**Figure 6**) and their survival in nature during the night.

Despite its advantages, the method has limitations. It requires access to a thermocycler, which may not be readily available in all laboratories, although such equipment has been present in laboratories for almost 40 years (33). Moreover, the cells are assessed under non-photosynthetic conditions, potentially limiting the relevance of the results to light-dependent processes. Another important consideration is that this assay specifically evaluates temperature tolerance rather than optimal growth temperatures. Although optimal growth temperatures are relevant, precise temperature control is often impractical in large-scale industrial algae cultivation, making tolerance a more critical factor to assess. The simplicity and efficiency of this assay provide a practical means to rapidly identify temperature-tolerant strains and effectively narrow down candidate pools for subsequent detailed analysis.

Future research should investigate the influence of incubation time on the screening results and the relationship between temperature tolerance and optimal growth temperatures. Building on these findings, this method could be employed to develop algae strains with enhanced temperature tolerance, improving their resilience and utility in diverse biotechnological applications.

### Light Tolerance

Light is the primary energy source driving microalgal photosynthesis, but it can also become a potent stressor when in excess or when fluctuating rapidly. Each algal strain has an optimal light intensity range for maximal photosynthetic productivity. Under low light, cells are limited by energy availability; under extremely high light, they risk photodamage to the photosynthetic apparatus. When photons saturate the light-harvesting capacity, the surplus energy can generate reactive oxygen species and cause oxidative damage to pigments and the proteins of Photosystem II, leading to a decline in photosynthetic efficiency known as photoinhibition (34).

Natural environments and open cultivation systems present microalgae with highly variable light conditions. In shallow ponds or bioreactors, cells circulate between the surface (full sunlight) and deeper layers (shade), experiencing light flashes and dark periods within seconds. Cloud cover, day/night cycles, and seasonal changes further modulate light intensity. Many microalgae can acclimate to different average light levels (by adjusting pigment content and photosynthetic unit size (35). For outdoor cultivation, high light tolerance is especially important: a strain should endure bright midday sun without severe photoinhibition. In open pond systems and other algae cultivation setups, light intensities can exceed 2000 µE/m²/s for extended periods (36). Some cultivation strategies (like using inclined panels or introducing light shading) can moderate intensity, but ultimately, an inherently light-tolerant strain is advantageous (18).

The results demonstrated the effectiveness of the light tolerance assay in distinguishing among *Chlamydomonas* strains based on their ability to tolerate varying light intensities (**Figure 8**). Cpa showed the highest light tolerance, with colonies appearing in regions with intensities exceeding >1200 µE/m²s, followed by moderate tolerance in Cin, while Cre displayed the lowest tolerance. The competitive growth and recovery of strains from mixed cultures further validated the method, with Cpa and Cin outperforming Cre under high-light conditions. This method represents a significant innovation in light tolerance screening for algae, requiring minimal specialized equipment besides strong LED light sources, which are widely available.

The light tolerance assay leverages the conical diffusion pattern of non-collimated light from LEDs to generate a gradient of incident intensities across a plate. A linear gradient is formed by positioning plates slightly to the center of the light source, enabling the effective screening of light tolerance across a range of intensities, varying the plate distance from the light source.

Additionally, this method facilitates the development of algae strains resilient to high light intensities, improving productivity in outdoor cultivation systems. This has already been demonstrated with a bred strain of Cpa that exhibited superior performance under high-light conditions (18).

Despite its strengths, the method has limitations. It requires the ability of strains to grow on agar plates and evaluates tolerance only for a fixed exposure time, rather than determining optimal light conditions. Furthermore, the study tested only three species, limiting the generalizability of the findings, though the method is likely applicable to other algae species. Future studies should investigate whether a similar setup could be adapted to determine optimal growth light intensities through image analysis of growth regions on plates or by combining multiple stressors in a single assay. Building on these results, further work could focus on improving strain tolerance to high light and assessing their performance in real-world cultivation systems, paving the way for enhanced productivity in algae-based biotechnology.

## Conclusion

This study highlights the development and validation of low-cost, accessible screening methods to evaluate the tolerance of algae strains to extreme environmental conditions, including pH, salinity, temperature, and light. By leveraging simple gradient-based systems and selective recovery assays, we successfully demonstrated the ability to distinguish strain-specific stress tolerance traits under controlled conditions. These methods proved effective in identifying individual strain responses and competitive co-culture settings, enabling the selection of the most resilient strains.

Each assay addressed critical environmental factors that directly influence algae growth and adaptability, such as pH, salinity, temperature, or light tolerance. The stability and reproducibility of these methods, combined with their ease of implementation, make them practical tools for laboratories worldwide, including those with limited resources. Importantly, these approaches are not limited to algae but can be adapted to other microorganisms, broadening their applicability in microbial ecology, biotechnology, and industrial strain selection. While each method demonstrated robustness and practical utility, limitations such as the need for solid media growth or fixed exposure times highlight areas for future refinement. Expanding the range of tested species, improving gradient dynamics, and incorporating advanced imaging and analysis techniques could further enhance the utility and precision of these assays. Overall, these methodologies provide a versatile and scalable platform for exploring the stress tolerance of algae and other microorganisms. Their ability to identify and improve traits critical for biotechnological and ecological applications ensures their relevance in advancing sustainable solutions for bioenergy, aquaculture, and environmental management challenges.

## Material and Methods

### Strain Cultivation and Media Preparation

#### Strains

Three Chlamydomonas species were used in this study: *Chlamydomonas pacifica* (Cpa, strain ID **CC-5697**), *Chlamydomonas incerta* (Cin, strain ID **CC-3871**), and the reference strain *Chlamydomonas reinhardtii* **CC-620** (mt^+^), available at the Chlamydomonas collection. All strains were maintained on solid Tris–Acetate–Phosphate (TAP) medium and in liquid culture under controlled conditions. Cultures were grown at 25 °C under continuous illumination (approximately 60 µmol photons m^−2^s^−1^) with shaking at 120 rpm, unless otherwise noted. For photoautotrophic growth experiments, cells were transferred to High Salt Minimal (HSM) medium (Sueoka’s medium), which lacks acetate and with NaNO_3_ as nitrogen source in equivalent amounts.

#### Media Composition

TAP medium (pH ∼7.0) consisted of a Tris base buffer with acetate as a carbon source (20 mM Tris, 9.3 mM NH_4_Cl, 1mM K_2_HPO_4_ 80 μM MgSO_4_, 68 μM CaCl_2_, trace elements (37), and 17 mM acetate). HSM medium (pH ∼6.8) is a minimal medium without organic carbon, containing the same major salts and trace elements as TAP but with NaNO3 as nitrogen source. 9.3 mM NaNO_3_, 14 mM K_2_HPO_4_ 80 μM MgSO_4_, 68 μM CaCl_2_, Kropat’s trace elements. Solid media were prepared by adding 1.5% (w/v) agar to the respective media. Starter cultures of each strain were grown to the stationary phase in TAP broth before inoculation into experimental assays described below.

### Vector Construction and Strain Engineering

#### Plasmid Design

All vectors were constructed using the pBlueScript II KS+ (pBSII) backbone. Vectors pJP32PHL7 and pAH04 (38,39) utilize the nuclear AR1 (HSP70/RBCS2) promoter to drive gene expression. The *ble* gene is the selectable marker in both constructs, linked to the respective genes of interest via a foot-and-mouth-disease virus 2A (FMDV-2A) self-cleaving peptide. Specifically, in the pJP32PHL7 vector, *ble* is fused with the FMDV-2A peptide and includes the signal peptide SP7 to facilitate secretion. In contrast, the pAH04 vector does not include a signal peptide sequence.

The pJPCHx1 vector was assembled utilizing genetic elements derived from the recently sequenced *Chlamydomonas pacifica* genome (9) and is thoroughly described in Molino et al., 2024 (18). Notably, this vector incorporates a hygromycin resistance marker, which was crucial for selective screening assays in this study.

All vector fragments were assembled using the NEBuilder® HiFi DNA Assembly kit (New England Biolabs, Ipswich, MA, USA), following the manufacturer’s recommended protocol. To facilitate linearization, each vector includes restriction enzyme sites flanking the expression cassette (XbaI at the 5’ end and KpnI at the 3’ end). Restriction enzymes were acquired from New England Biolabs (Ipswich, MA, USA). Final sequences for all vectors are publicly available on Zenodo (https://zenodo.org/records/15357688), and detailed vector maps are provided in **Supplementary Figure 7**.

#### Transformation and Strain Generation

The constructed plasmids were introduced into the appropriate Chlamydomonas strains to create engineered lines for competition experiments. *The Chlamydomnas strains* were transformed with the nuclear vectors by an electroporation method (38,39). Briefly, mid-log cells (approximately 3-6 x 10^6^ cells) were mixed with 0.5-1.0 µg of plasmid DNA (linearized with XbaI/KpnI) and electroporated in a time constant protocol at 800V, for 20 us in a 4 mm electroporation cuvette. *Cre* CC-620 was transformed to introduce the pJPCHx1 construct, while *Cin* received the pAH04 and *Cpa* pJP32. After DNA delivery, cells were allowed to recover overnight in TAP medium, then plated onto selective TAP agar. Transformants carrying pJP32PHL7 or pAH04 were selected on TAP + Zeocin (15 µg/mL), while those with pJPCHx1 were selected on TAP + hygromycin B (30 µg/mL). Single colonies were picked after 7–14 days and re-streaked on the same selective media to obtain clonal strains. Successful integration of transgenes at *Cre* and *Cin* was verified by PCR of genomic DNA (40), and *Cpa* was confirmed by clearing zones around colonies in a plastic polydispersion plate in agar. These engineered strains (Zeocin-resistant and Hygromycin-resistant) were used in subsequent gradient and competition assays.

### pH Gradient Plate Assay

A solid-phase gradient assay was developed to evaluate growth tolerance across a continuous pH range. **Gradient Generation:** HSM agar plates (90 mm Petri dishes) were prepared with the addition of a pH indicator dye (Fisher Chemical Universal pH Indicator System), final concentration ∼2% v/v) to visualize pH changes in the pH characterization experiments, and was absent for experiments with cells. Once the agar solidified, a strong base and acid were applied at opposite edges of the plate to create the gradient. Specifically, 120 µL of 10 M KOH was pipetted onto the surface at one side of the agar, and 60 µL of 3.3 M H_3_PO_4_ was applied to the opposite side (**Figure 1, Panel A, B**). The solutions were allowed to diffuse into the agar for 24-48 hours at room temperature with the lid closed. The Universal pH indicator facilitated visualization of the resulting pH gradient, shifting from yellow/orange in acidic regions to cyan/blue in basic regions. By 2 hours post-application, a stable color across the plate confirmed the establishment of pH regions (below pH 4 to above pH 10 spanning the plate).

#### Cell Seeding and Incubation

After the pH gradient was formed, each strain was inoculated onto the plate to assess its viable pH range. For single-strain tolerance tests, ∼10^6^ cells of a given strain were evenly spread onto the gradient plate surface using a sterile plastic loop, taking care to distribute cells across the entire pH range. Plates were incubated under standard light conditions (continuous light, ∼50 µmol/m2s1) at 25 °C for 7–10 days. Growth was evident as lawn or colony formation in regions of the plate where the local pH supported viability. The pH value at the edge of visible growth was noted for each strain as an estimate of its tolerance limits.

#### Buffer Variations

To examine the effect of medium buffering capacity on the pH gradient and cell survival, we prepared HSM agar with different buffering systems in separate experiments. In one variant, the standard HSM phosphate buffer (KH_2_PO_4_/K_2_HPO_4_, ∼2 mM) was replaced with a 50 mM citrate buffer (pH 7.0) before autoclaving. In another, a combined buffer system of HSM + 25 mM citrate (pH 7.0) was used. Gradient plates were formed as described above on these buffered media (using KOH and H_3_PO_4_ at the same concentrations). The time required for gradient formation and the shape of the pH profile were recorded. We observed that different buffering (citrate and combined) changes the pH range profile, as expected due to different pKa and buffer capacity. These plates were likewise incubated, and a time-lapse video was recorded to observe the pH variation per region in the plate over time (**Video 1**).

### NaCl Gradient Plate Assay

To assess salinity tolerance, we developed an analogous gradient plate method for sodium chloride. **Gradient Formation:** HSM agar plates were modified by incorporating a high-salt agarose section on one side of the plate. First, a base layer of normal HSM agar (1.5% agar) was poured to cover approximately the Petri dish. After solidification, we sliced out approximately 10% of the plate area in one corner and filled it with a 1.5% low-melt agarose gel containing 5 M NaCl solution. This high-salt agarose section was allowed to solidify, forming a contiguous gel with the rest of the plate. The plate was then incubated at room temperature for 2 hours to let the gel solidify. Over time, a concentration gradient of NaCl was established across the plate, ranging from near 5 M at the salt side to ∼0 M at the far side. Although not directly visible, the gradient was inferred by the diffusion time and later confirmed by biological response (growth inhibition patterns).

#### Cell Seeding and Growth

Cells of each strain were uniformly spread on the surface of the gradient plate (∼10^6^ cells per plate, as for the pH assay). The plates were incubated at 25 °C under constant light for 7–10 days to allow colonies to develop in permissive regions. Salt-sensitive strains only formed colonies on the low-salt half of the plate, whereas more salt-tolerant strains grew further toward the high-salt side. The distance of colony growth into the high-salt region was used as a qualitative measure of salt tolerance.

#### Liquid Assay for Validation

To quantitatively validate the salt tolerance differences observed on gradient plates, we performed a liquid growth assay across discrete salt concentrations. Cells were inoculated into 96-well microplates containing 200 µL HSM medium with NaCl concentrations of 0, 0.5, 1.0, 1.5, and 2.0 M (in triplicate wells for each strain and concentration). The microplate cultures were grown for 7 days under continuous light (∼60 µmol m^−2^s^−1^) at 25 °C without shaking. Growth was monitored by measuring in vivo chlorophyll fluorescence (excitation 440 nm, emission 680 nm) using a plate reader. Fluorescence values (relative fluorescence units, RFU) were recorded as an indicator of biomass. To account for background and pathlength differences, readings were normalized to the 0 M control (set as 100%). These data provided a quantitative salt tolerance curve for each strain.

### Temperature Gradient Assay Using a Thermocycler

Thermotolerance of the strains was evaluated by incubating cells across a range of temperatures simultaneously using a thermal cycler gradient block. **Setup:** Aliquots of stationary phase cultures (100 µL each) were dispensed into the wells of an 8×12 PCR plate (96-well format). The PCR plate was sealed with a clear film to prevent evaporation and placed in a PCR machine with gradient capability. A temperature gradient from 34 °C (min) to 42 °C (max) was applied across the plate for cell characterization and 35 °C (min) to 40 °C (max) for cell mixtures. The plate was incubated in the dark (no light) for 24 hours under these set temperatures, allowing each subset of wells to experience a constant, distinct temperature. During this incubation, cells remained in stationary culture (no shaking).

#### Post-Incubation Spotting

To assess survival and growth after exposure, we performed a spotting assay following the temperature treatment. Immediately after the incubation period, 5 µL from each well was spotted onto fresh TAP agar plates (non-selective) in an ordered grid corresponding to the original temperature gradient. Four replicate spots were plated for each temperature condition to ensure statistical reliability. The spotted plates were then incubated under standard growth conditions (25 °C, continuous low light) for 5–7 days. Viable cells from the temperature-treated aliquots formed visible microcolonies at the spot locations. We recorded the highest temperatures at which growth occurred for each strain. Strains incapable of surviving extreme temperatures failed to produce colonies from those spots. Using this method, we determined the thermal tolerance range of each Chlamydomonas strain.

### Light Gradient Assay

A light intensity gradient was employed to test the photo-tolerance of the strains, especially under extreme irradiance. **Gradient Generation:** Cells were plated on solid minimal medium (HSM agar, 1.5% agar) and exposed to a spatial gradient of light intensity (**Figure 8, Panel A and B**). To create the gradient, the plates were placed at a distance of 9 cm above a 1000W Mastiff GrowL® LED grow light, which emits both blue and red wavelengths to mimic high-intensity illumination. To establish a gradient of light intensity, the plates were arranged such that some areas were directly illuminated by the LEDs, while other areas extended beyond direct exposure. We also added a filter with different light transmittance. The filter was generated using a laser printer, with different shades of black added to a white paper with a basis weight of 75 g/m^2,^ generating values between 40 µE m^−2^s^−1^ (under the darkest end) and 500 µE m^−2^s^−1^ (under the least dark end). The other half of the plate was left uncovered, receiving direct illumination from the light source. The illumination delivered up to ∼3000 µE m^−2^s^−1^ to the fully exposed side.

#### Exposure and Recovery

*Chlamydomonas* cells were pre-plated uniformly on the HSM agar (approximately 10^6^ cells spread over the plate) prior to applying the light stress. The plate was then immediately subjected to the gradient light exposure for 24 hours continuous duration. During this period, the plate was maintained at 25 °C, and evaporation was minimized by sealing the edges with Parafilm. After 24 hours of high-intensity light treatment, the plate was removed from the light source and the neutral-density filter was taken off. The plate was then transferred to standard growth conditions (low light ∼50 µE m^−2^s^−1^, 25 °C) for a recovery period of 7-10 days. Surviving cells grew into colonies during the recovery phase. Because cells in different regions of the plate had received different light doses, the pattern of colony formation reflected the tolerance limit. Regions that had been exposed to ∼500 µE m^−2^s^−1^ or less (under the filter) generally showed robust colony growth, whereas the areas that experienced >2000 µE m^−2^s^−1^ of direct light had few or no surviving colonies (depending on the strain). The maximum light intensity that still allowed colony formation was noted for each strain, indicating its photo-tolerance threshold.

### Competitive Growth and Selective Recovery Assays

Competitive co-culture assays were performed on each gradient platform to directly compare the fitness of different strains under stressful conditions and validate the method’s capacity to discern among strains (**Supplementary Figure 6**). In these experiments, two strains—each marked with a distinct antibiotic resistance—were mixed and exposed to the environmental gradient, and their relative survival was measured by selective plating. **Mixing Strategy:** Competing strains were prepared by growing each separately to mid-log phase in TAP liquid medium, then washing and resuspending in fresh medium. Pairs of strains were mixed at defined ratios immediately before application to the gradient assay. Two mixing ratios were tested: a 1:1 ratio (equal cell numbers of each strain, ∼5×10^5^ cells of each), and a 1:100 ratio (minority:majority) to simulate a scenario where one strain was initially rare. In the 1:100 mixes, ∼5×10^4^ cells of the minority strain were mixed with ∼5×10^6^ cells of the majority strain. For example, *Cpa* (Zeo^R^) and *Cre* (Hyg^R^) were mixed in equal numbers for one set, while in another set *Cpa* was the minority against a majority of *Cre*.

#### Exposure on Gradient Assays

The mixed cell populations were introduced to the gradient assays in the same manner as single-strain tests. For pH and salt gradient plates, the mixed culture was spread uniformly across the plate surface. For the thermocycler temperature assay, equal aliquots of the mixed culture were added to each well of the PCR plate (in place of single-strain inoculum). For the light gradient, the two strains were co-spread on the agar plate. In all cases, the mixed cultures were concurrently subjected to the environmental gradient challenge. After the exposure period (and recovery, if applicable, as in the light assay), cells from the assay were harvested for selective plating. In the case of gradient plates (pH, NaCl), the population of cells in the transition zone was collected. The cell suspension was thoroughly mixed to ensure homogeneity, then appropriate dilutions were prepared. For the temperature gradient, cells from each temperature well were spotted on TAP, TAP/Zeo, and TAP/Hyg plates to evaluate the identity of the surviving strain. Individual colonies were picked and grown in 96-well plates in the transition zone for the light experiment and plated on TAP, TAP/Zeo, and TAP/Hyg plates to evaluate their identity.

#### Selective Plating

To determine the surviving fraction of each strain in the mix, aliquots of the harvested cell suspensions were plated on two types of selective media: TAP + Zeocin (15 µg/mL) to recover the strain carrying the ble (Zeo^R^) marker, and TAP + hygromycin B (30 µg/mL) to recover the strain with the Hyg^R^ marker. For each treatment, 100 µL of a suitable dilution was spread on three replicate plates for each antibiotic selection. Plates were incubated for 10–14 days at 25 °C in moderate light until colonies formed. Colony counts on Zeo plates versus Hygro plates reflected the number of survivors of each strain.

### Data Collection and Analysis

Experiments were conducted with four independent replicates to ensure reproducibility. **Imaging and Colony Quantification:** Gradient plates (pH, salt, light) were imaged at the end of the incubation/recovery period using a Canon camera and EOS utility Software. The pH gradient plates after incubation (to record growth). Colony formation patterns were documented, and the images were later analyzed to observe the relative tolerance among strains. For competitive assays, selective plates were imaged after colony development. Colony counting was performed manually for each plate; in cases of high colony density, counts were assisted by the OpenCFU software (open-source colony counting tool) to improve accuracy (41). For microplate-based fluorescence measurements (salt tolerance validation), data were exported from the plate reader software as raw fluorescence units. Background fluorescence (medium-only wells) was subtracted, and values were normalized to the no-stress control for each strain.

#### Normalization and Statistical Analysis

For competition outcomes, the proportion of surviving cells of each strain was calculated as the number of colonies on the specific selective medium. Statistical analyses were performed using R statistical software. Differences in tolerance between strains were evaluated by one-way ANOVA followed by Tukey’s post hoc test for multiple comparisons. A significance threshold of *p* < 0.05 was applied for all tests. Exact *p*-values for key comparisons (strain differences at tolerance limits, competition outcomes) are reported in the Results section. All data are presented as mean ± standard deviation of replicates.

## Author contributions

**BS:** Investigation, Methodology, Writing – review & editing

**KK:** Investigation, Methodology, Formal analysis, Visualization, Writing – original draft, Writing – review & editing

**LM**: Investigation, Methodology, Writing – review & editing

**AG**: Investigation, Writing – review & editing

**EES:** Investigation, Methodology, Writing – review & editing

**SM:** contributed to drafting and revising the original manuscript and secured funding for the research.

**JVDM**: Conceptualization, Data curation, Formal analysis, Investigation, Methodology, Visualization, Writing – original draft, Writing – review & editing

