## Supplementary Figures for "Low-Cost Screening of Algae for Extreme Tolerance to pH, Temperature, Salinity, and Light"

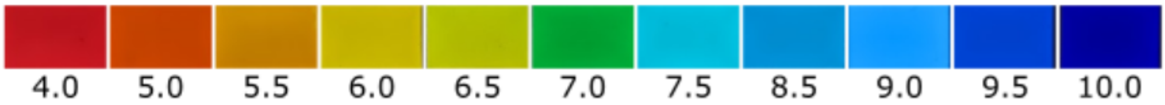


**Supplementary Figure 1: Fisher Chemical™ Universal pH Indicator System color scale for pH gradient visualization.** The image displays the color spectrum produced by the Fisher Chemical™ Universal pH Indicator System (Catalog number: SI60-500), which was used to visualize pH gradients in experimental setups. The indicator demonstrates a clear progression of colors across the pH range, transitioning from acidic to basic conditions. At pH 4.0, the indicator appears red, gradually shifting to orange and yellow between pH 5.0 and pH 6.5, representing weakly acidic conditions. Near-neutral pH values, around pH 7.0, produce a green coloration, while slightly basic values (pH 7.5–8.5) transition to cyan and light blue. Strongly basic conditions at pH 9.0–10.0 are indicated by darker blue shades.

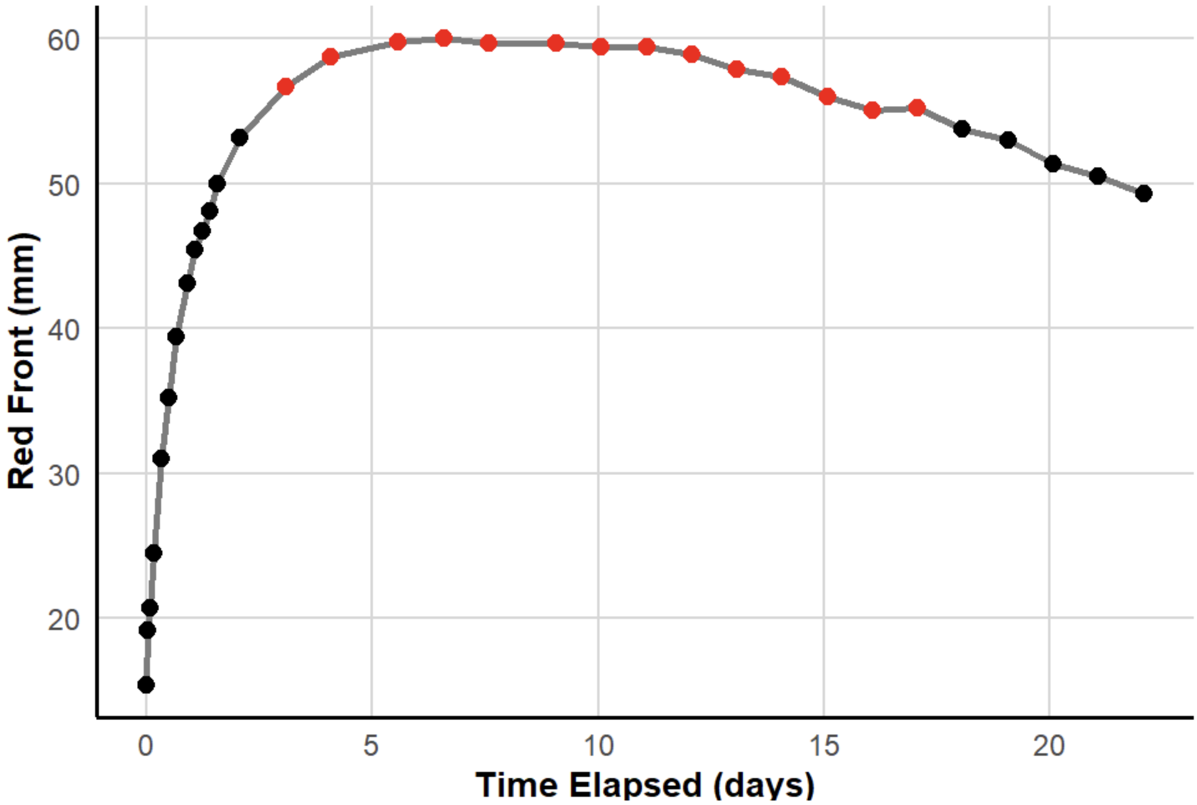


**Supplementary Figure 2: The position of the red front over time in the pH gradient plate was visualized using a pH indicator.** The graph illustrates the progression of the red front, corresponding to the acidic region (pH ~4.0–5.0), on a pH gradient plate over time. The x-axis represents the time elapsed (in days), while the y-axis shows the position of the red front (in millimeters) from the acid plate edge. The red data points indicate positions within 10% of the maximum value, marking a stable range between days 3 and 17. Initially, the red front expands rapidly, reaching its maximum position (~60 mm) within the first 5 days, after which it stabilizes. After 12 days, a gradual decline in the red front position is observed, indicating changes in the pH gradient, likely due to diffusion dynamics or buffer interactions within the plate. This trend demonstrates the temporal stability of the pH gradient.

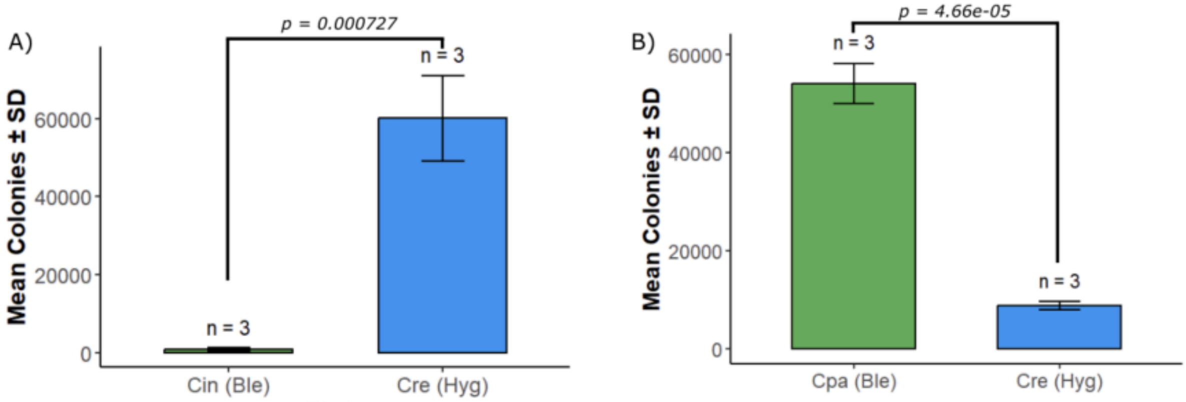


**Supplementary Figure 3: Colony recovery and selection of *Chlamydomonas* strains from the transition zones of pH gradient plates on the higher pH zone. (A)** Recovery of colonies from the transition zone of the *Chlamydomonas incerta* (Cin, Ble) and *Chlamydomonas reinhardtii* (Cre, Hyg) co-culture. Colonies appearing in the transition zone were scraped, resuspended in fresh media, and plated on hygromycin- (selecting for Cre) or zeocin- (selecting for Cin) containing plates. Colony counts revealed a significantly higher recovery of Cre colonies than Cin colonies, indicating more remarkable survival and competitive fitness of Cre in the transition zone. Mean colony counts (±SD) are presented for three replicate plates (n = 3). **(B)** Recovery of colonies from the transition zone of the *Chlamydomonas pacific* (Cpa, Ble) and *Chlamydomonas reinhardtii* (Cre, Hyg) co-culture. Colonies were similarly scraped, resuspended, and plated on hygromycin- (selecting for Cre) or zeocin- (selecting for Cpa) containing plates. In this pairing, Cpa demonstrated significantly higher colony recovery, reflecting its superior competitive ability in the basic pH region of the transition zone. Mean colony counts (±SD) are shown for three replicate plates (n = 3).

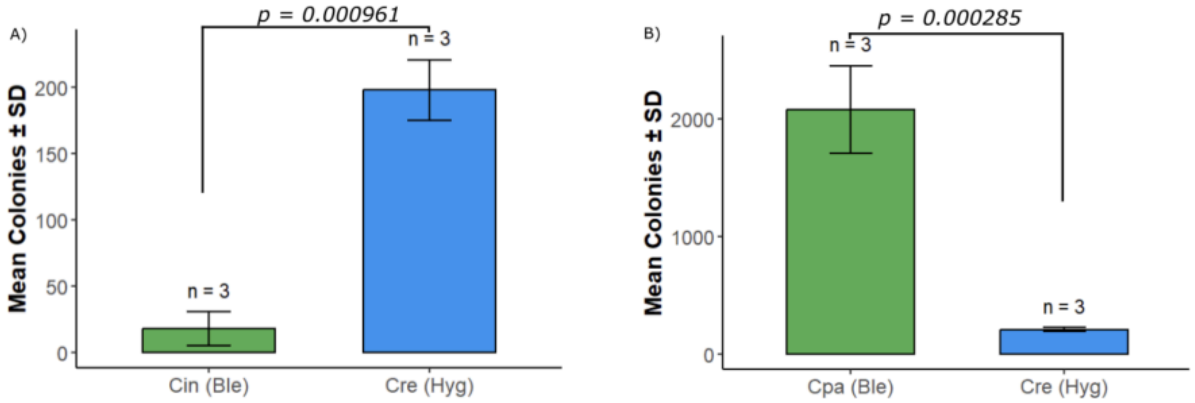


**Supplementary Figure 4: Colony recovery and selection of recombinant *Chlamydomonas* strains from the transition zones of NaCl gradient plates. (A)** Recovery of colonies from the transition zone of the *Chlamydomonas incerta* (Cin, Ble) and *Chlamydomonas reinhardtii* (Cre, Hyg) co-culture. Colonies were similarly scraped, resuspended, and plated on hygromycin- and zeocin-containing plates. Colony counts indicate that Cre strongly outcompetes Cin, as evidenced by significantly higher recovery of Cre colonies on hygromycin plates than Cin colonies on zeocin plates. Mean colony counts (±SD) are shown for three replicate plates (n = 3). **(B)** Recovery of colonies from the transition zone of the *Chlamydomonas pacific* (Cpa, Ble) and *Chlamydomonas reinhardtii* (Cre, Hyg) co-culture. Colonies appearing in the transition zone were scraped, resuspended in fresh media, and plated on hygromycin-containing plates (selecting for Cre) and zeocin-containing plates (selecting for Cpa). Colony counts reveal that Cpa significantly outcompetes Cre in the transition zone, as indicated by higher mean colony recovery on zeocin plates compared to hygromycin plates. Mean colony counts (±SD) are presented for three replicate plates (n = 3).

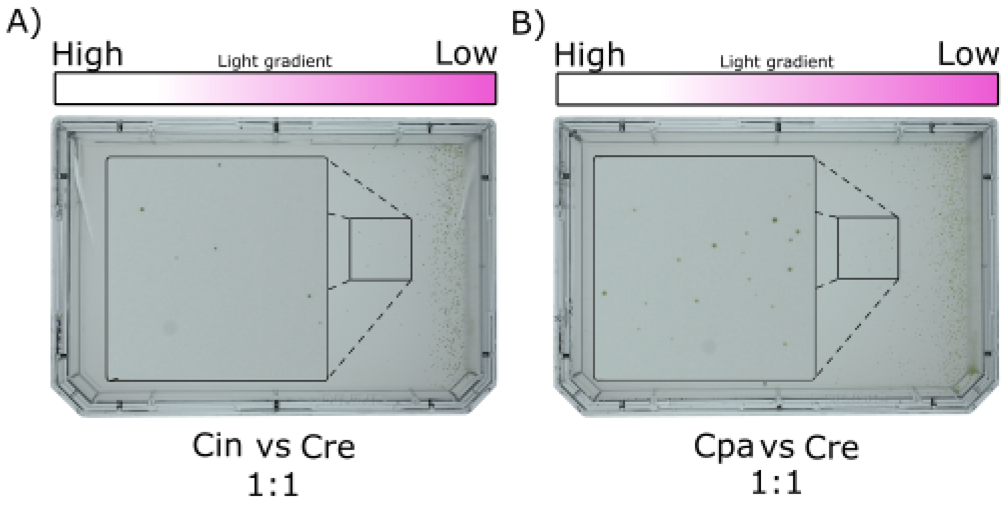


**Supplementary Figure 5: Competitive growth of *Chlamydomonas* strain mixtures under a light gradient. (A)** Competitive growth of *Chlamydomonas incerta* (Cin) and

*Chlamydomonas reinhardtii* (Cre) in a 1:1 mixture under a light intensity gradient. Cin harbors the vector pAH04mCherry, conferring resistance to bleomycin, while Cre harbors pJPCHx1_mVenus, conferring resistance to hygromycin. The light gradient ranged from high intensity (~3000 μE/m²/s) on the left to low intensity (~100 μE/m²/s) on the right. The dashed box highlights a magnified region showing the differential growth of the two strains, with Cre exhibiting better survival in regions of higher light intensity compared to Cin. **(B)** Competitive growth of *Chlamydomonas pacific* (Cpa) and *Chlamydomonas reinhardtii* (Cre) in a 1:1 mixture under the same light gradient. Cpa harbors the vector pJP32PHL7, conferring resistance to bleomycin, while Cre harbors pJPCHx1_mVenus, conferring resistance to hygromycin. The dashed box highlights a magnified region of the plate, demonstrating colonies growing in the high-light regions.

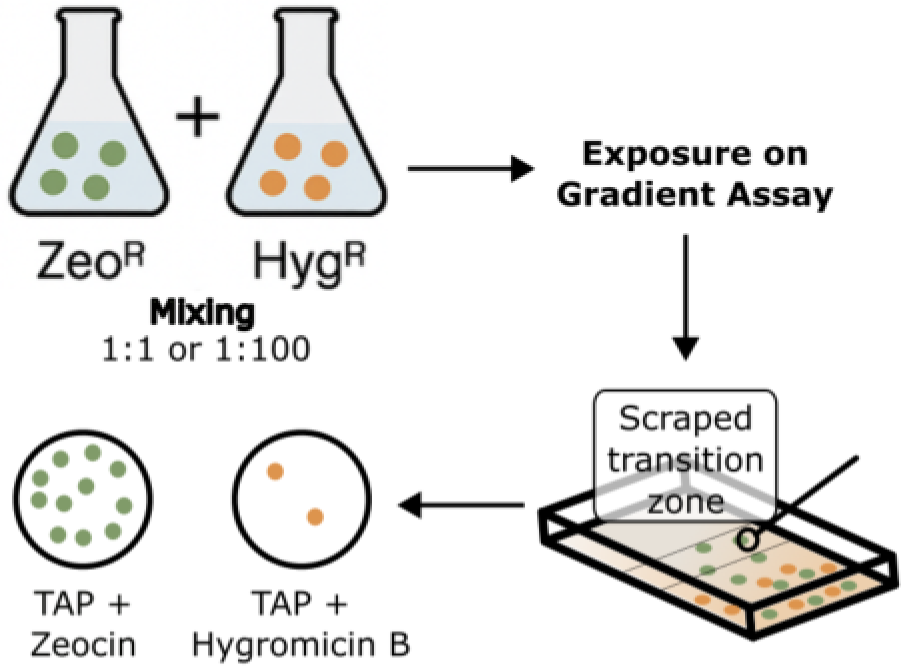


**Supplementary Figure 6: Schematic representation of competitive co-culture assays under environmental gradients.** Strains carrying distinct antibiotic resistance markers (Zeo$^R$ and Hyg$^R$) were grown separately to mid-log phase, mixed at defined ratios (1:1 or 1:100), and introduced onto environmental gradient assays (e.g., pH, salinity). Following exposure, cells were harvested from the transition zone of the gradient, resuspended, and plated onto selective media containing Zeocin or Hygromycin B to assess the relative survival of each strain. This approach enables the direct comparison of strain fitness under stress and validates the screening platform’s discriminatory capacity.

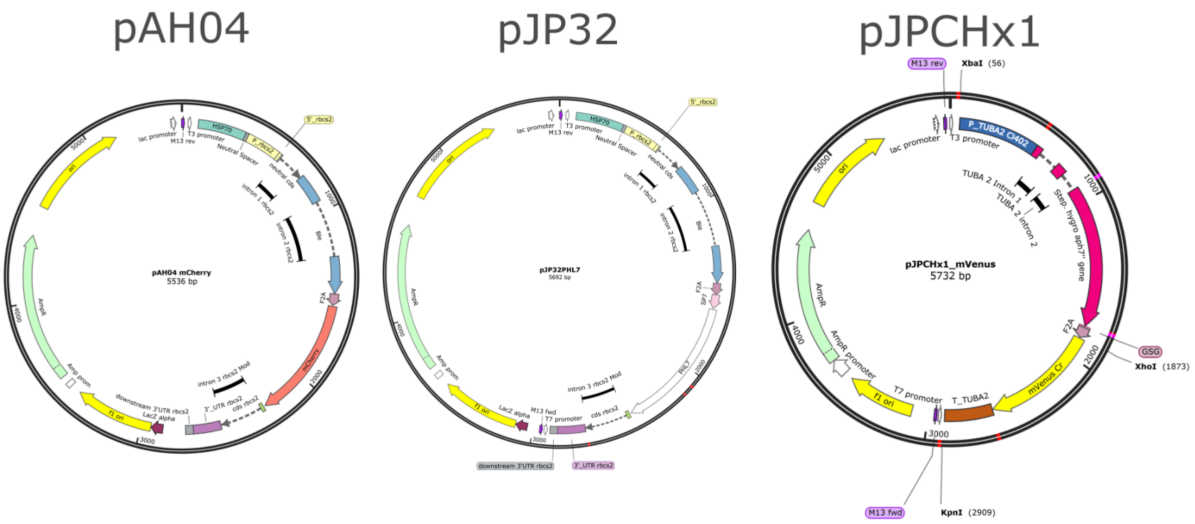


**Supplementary Figure 7: Schematic representation of the vectors used.** The whole sequence can be found at Zenodo. Vectors are available at the Chlamydomonas collection.

### Video

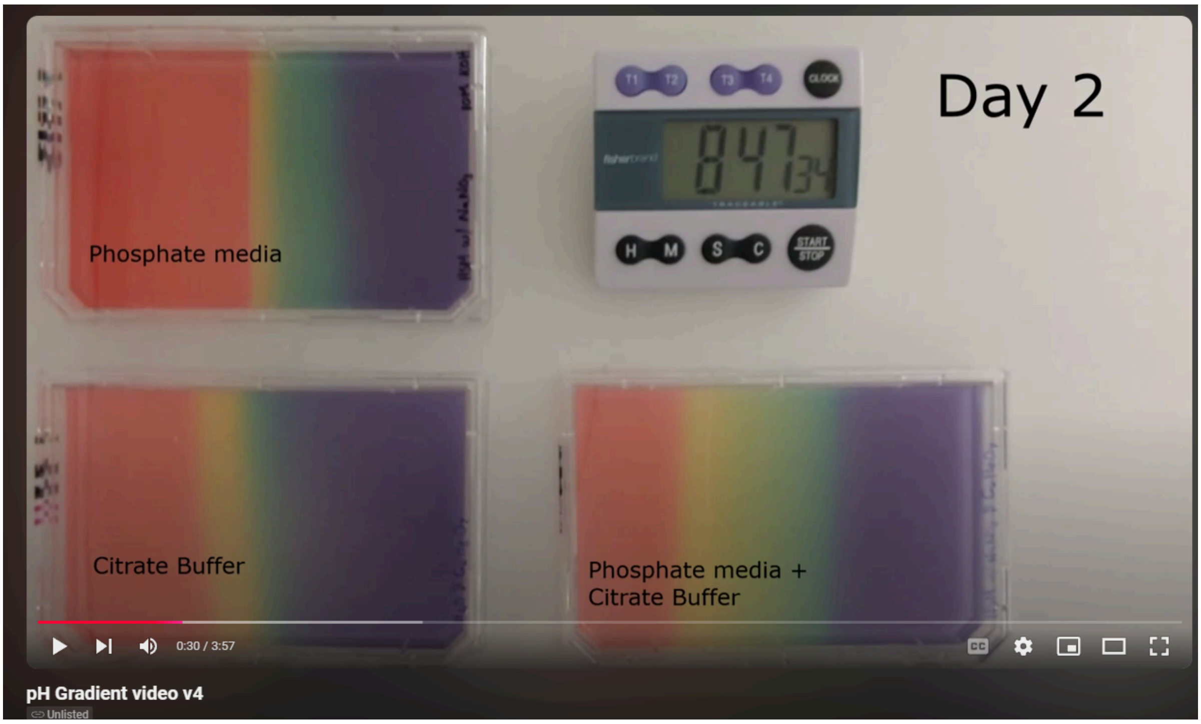


**Supplementary Video 1: pH Gradient Formation in Different Buffer Systems Over Two Weeks.** This video demonstrates the formation and stabilization of pH gradients on agar plates containing three different buffer systems—phosphate media (HSM), citrate buffer (100mM), and a combination of phosphate media (HSM) with citrate buffer—over a period of two weeks. A fixed volume of 160uL of 10 M KOH (base) was applied to one corner of each plate, while an 80uL of 3.3 M phosphoric acid (acid) was applied to the opposite corner. The pH gradients were visualized using a universal pH indicator solution, producing color transitions corresponding to pH variations (red = acidic, green = neutral, purple = basic). This gradient remained stable throughout the two-week period. The video highlights the time-dependent dynamics of pH gradient formation and stabilization across the different buffer systems. Citrate has three pKa, 3.13, 4.76, and 6.40. Phosphate also has three pKa in various regions: 2.16, 7.21, and 12.32 (1). The difference in pKa explains the difference in the pattern observed with the pH indicator. (Video 1)

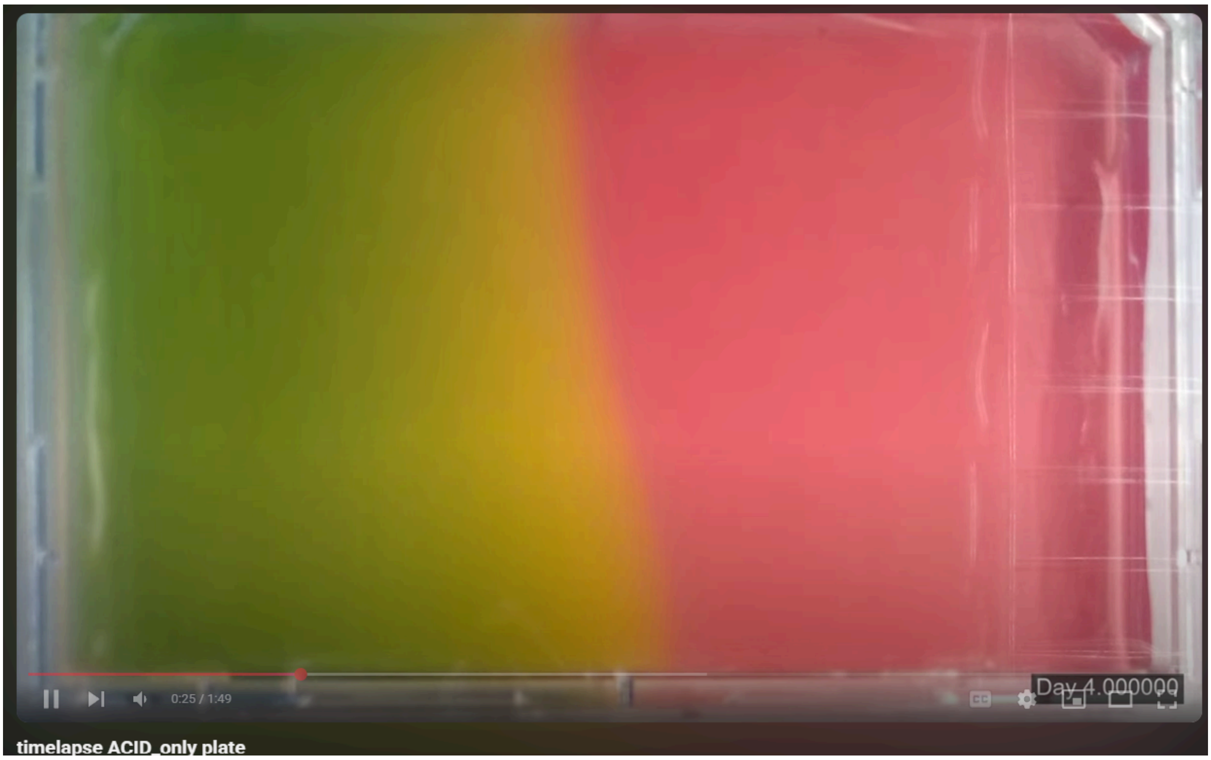


**Supplementary Video 2: Varying pH Gradient Formation in Buffer Systems Over Two Weeks with acid addition in the right corner.** This video demonstrates the formation and dynamic variation of pH on agar plates containing —phosphate media (HSM) —over a period of two weeks. A fixed volume of 80uL of 3.3 M phosphoric acid (acid) was applied to the right corner. The pH gradients were visualized using a universal pH indicator solution, producing color transitions corresponding to pH variations (red = acidic, green = neutral, purple = basic). This gradient remained unstable throughout the two-week period. The video highlights the time-dependent dynamics of pH gradient formation and variation across the plate. (Video 2)
